# The ITCC-P4 PDX platform of pediatric cancers for preclinical testing

**DOI:** 10.64898/2026.02.04.703023

**Authors:** Aniello Federico, Apurva Gopisetty, Didier Surdez, Yasmine Iddir, Robert J. Autry, Joshua Waterfall, Elnaz Saberi-Ansari, Charles Bobin, Stelly Ballet, Gaelle Pierron, Justyna Wierzbinska, Andreas Schlicker, Martin Sill, Richard Volckmann, Danny A. Zwijnenburg, Alan Mackay, Sakina Zaidi, Alexandra Saint-Charles, Norman Mack, Benjamin Schwalm, Lena Weiser, Ivo Buchhalter, Christopher Previti, Anna-Lisa Böttcher, Fatima Iradier, Eva-Maria Rief, David T.W. Jones, Olaf Witt, Frank Westermann, Till Milde, Angelika Eggert, Nicole Huebener, Johannes Schulte, Sara Colombetti, Louis Chesler, Heinrich Kovar, Jan-Henning Klusmann, Klaus-Michael Debatin, Simon Bomken, Christina Guttke, Petra Hamerlik, Maureen Hattersley, Michelle Garcia, Frédéric Colland, Ashley Strougo, Pablo M. Aviles, Michal Zapotocky, David Sumerauer, Johannes Gojo, Walter Berger, Daniela Lötsch-Gojo, Julia Schueler, Emily Girard, James M. Olson, Beat Schäfer, Marco Wachtel, Jan J. Molenaar, Biljana Dumevska, Elizabet Fernandez Potente, Chris Jones, Ángel M. Carcaboso, Emilie Indersie, Stefano Cairo, Katia Scotlandi, Maria Cristina Manara, Lorena Landuzzi, Angela Di Giannatale, Patrizia Gasparini, Massimo Moro, Dennis Gürgen, Jens Hoffmann, Maria Eugénia Marques da Costa, Birgit Geoerger, Olivier Delattre, David J. Shields, Hubert N. Caron, Gilles Vassal, Lou F. Stancato, Stefan M. Pfister, Natalie Jäger, Jan Koster, Marcel Kool, Gudrun Schleiermacher

## Abstract

Cancer is the leading cause of disease-related deaths among children in high-income countries. Tumor heterogeneity and lack of mechanism-of-action-based therapeutic options are key challenges to overcome in order to improve pediatric cancer patients’ survival. Here, we report the EU-IMI-2 funded public-private partnership “ITCC-Pediatric Preclinical Proof-of-Concept Platform” (ITCC-P4), which has built a large repertoire of patient-derived xenograft (PDX) models, representing all major solid pediatric cancer types, for *in vivo* drug testing. Three-hundred-fifty-three PDX models from diagnostic and relapsed pediatric cancers have been established and molecularly characterized, together with matched germline/tumor samples. As proof-of-concept, we present *in vivo* drug screening data in neuroblastoma and rhabdomyosarcoma models. PDX data, accessible at http://r2platform.com/itcc-p4, allow the selection of models based on oncogenic drivers and/or potential biomarkers for preclinical testing. Operated by a non-profit entity (www.itccp4.com), this sustainable platform aids academic and industrial researchers in developing and prioritizing innovative therapies for pediatric cancer.

**Graphical abstract:** 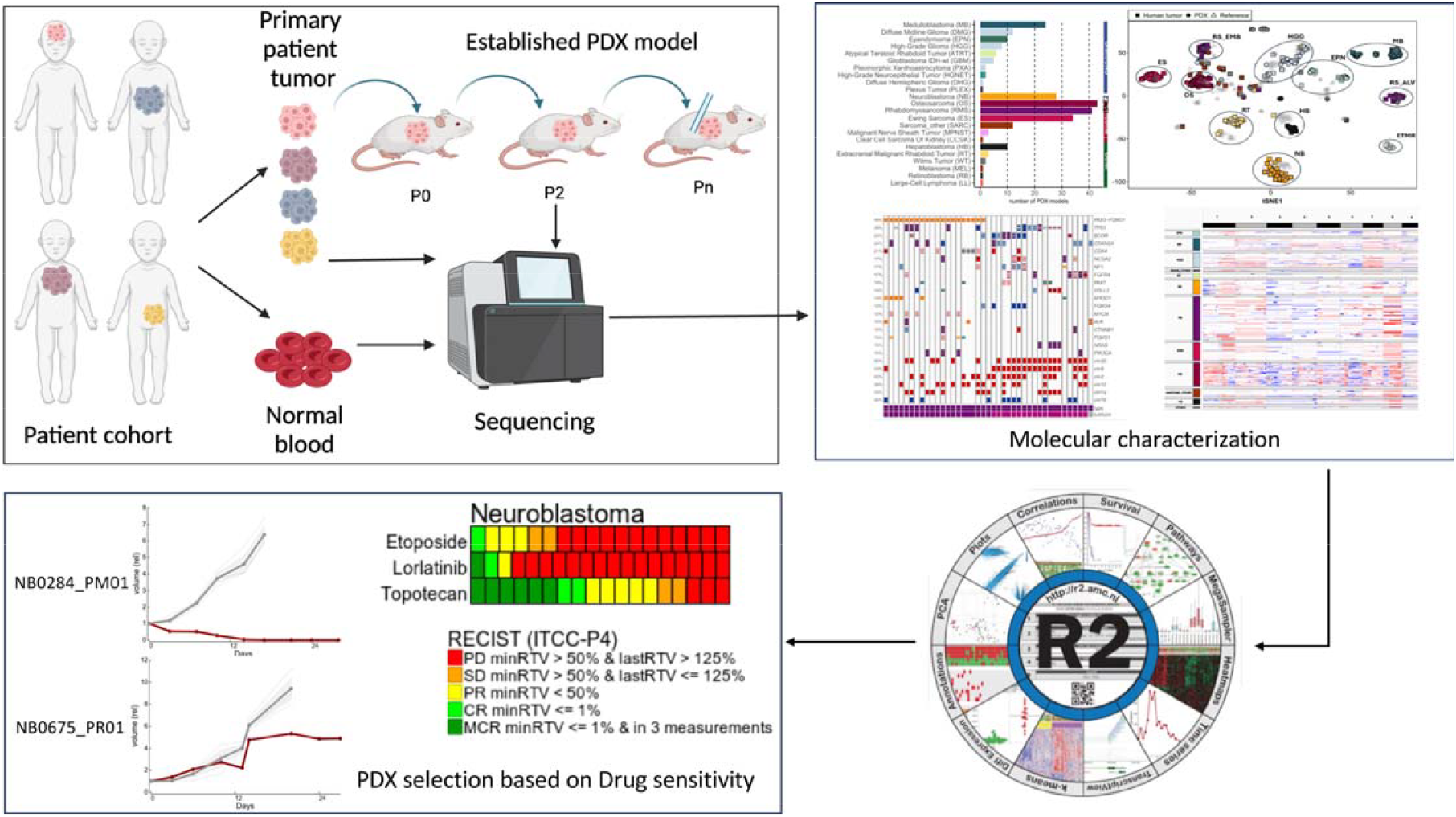

## Introduction

Cancer represents the leading cause of disease-related death in childhood in high-income countries. In pediatric cancers, their embryonal origins frequently contribute to their unique epidemiology and etiology. They are characterized by a relatively low mutational burden, high inter-tumoral heterogeneity and cellular plasticity. The most frequent pediatric cancers are acute leukemias and lymphomas, followed by central nervous system (CNS) tumors and solid extracranial tumors such as neuroblastoma, soft-tissue sarcoma, and bone sarcoma.

Despite many advances in pediatric cancer research, a lack of effective therapeutic options for patients with high-risk disease or treatment failure has hampered improvement in patients’ survival. Precision medicine programs such as INFORM, MAPPYACTS, iTHER, MATCH, and KiCS, reconfirm the general rarity of specific targetable molecular alterations, and only a subset of patients can benefit from identified targeted treatments (1–4). Furthermore, most childhood cancer survivors experience long-term treatment-related sequelae (5). Coordinated efforts and systematic, multi-disciplinary approaches are required to accelerate drug development and discover novel therapeutic approaches for children with cancer.

The development of new, effective childhood cancer drugs is hindered by limited market interest and increasingly stringent regulatory standards. However, initiatives such as the Research to Accelerate Cures and Equity (RACE) for Children Act highlight a significant regulatory shift towards prioritizing pediatric cancer treatments and fostering tailored therapies (6). To test novel therapies in clinically relevant models, patient-derived tumor xenografts (PDXs) have been broadly developed. PDXs tend to preserve the main molecular features of the original tumors (7–10) and response to treatment in PDX has been shown to correlate with that observed in patients (11,12). Hence, these models are currently regarded as the most predictive for evaluating new drugs for pediatric clinical trials and investigating treatment resistance mechanisms *in vivo* (13).

Large collections of molecularly well-characterized PDX models are currently available for a wide variety of adult cancers. These collections have supported the development of clinical trials through the identification and investigation of predictive biomarkers for stratified treatment strategies, the systematic and rigorous *in vivo* testing using single-mouse trial approaches, and the definition of subtype-specific drug responses (14). For pediatric tumors, molecularly characterized PDXs for CNS malignancies, neuroblastoma and sarcoma, as well as recurrent refractory pan-cancer cohorts, have also been established more recently (8,10,15–19). However, the inaccessibility of these models for commercial drug testing limits their utility in preclinical evaluations for the development of innovative drugs aimed at treating children with cancer.

Persistent challenges for pediatric PDX cohorts include a) collecting sufficient numbers of PDXs to cover the full spectrum of molecular subtypes for a given tumor entity and to cover the high intra- and inter-tumoral heterogeneity between and within subtypes; b) including models from different clinical phases, such as treatment-naïve and treatment-resistant models, to study the mechanisms underlying the response/resistance to prior therapies; c) fully integrating molecular data from PDXs with patient tumor and germline data, thus recapitulating the molecular landscape of these tumors; d) optimizing generation and application of these models in preclinical studies in standardized and harmonized workflows that adhere to the new regulatory requirements and bridge gaps between academic groups and industry; and e) enabling broad access to these PDX models for preclinical drug testing.

Based on these objectives, a public-private partnership comprised of academic and industry partners formed in 2017 in an initiative called “Innovative Therapies for Children with Cancer Paediatric Preclinical Proof-of-concept Platform” (ITCC-P4; itccp4.eu), which in 2023 transitioned into a non-profit company (www.itccp4.com). The main goal of the project was to build a sustainable platform of molecularly well-characterized PDX models representing the full spectrum of most malignant pediatric tumors for drug testing and preclinical prioritization of novel therapies for pediatric cancer patients. In addition to a large collection representative of the main pediatric tumor types, our PDX collection also includes many high-risk childhood cancer entities for which no appropriate preclinical models are currently available, as well as models generated from relapsed tumors. All models underwent comprehensive molecular characterization to identify potential biomarkers for predicting therapeutic response in patients. The availability of matching patient material (tumor and/or germline control) for most cases allowed a direct (somatic) molecular comparison between PDX and original patient tumors.

Here, we describe the first 353 fully established and extensively characterized PDX models that represent a diverse spectrum of pediatric malignancies, including 119 models (47%) from refractory or relapse disease. A subset of serial PDX models generated from tumors collected at different disease stages from the same patients enabled the investigation of molecular changes driving tumor progression and treatment responses. Our integrative, multi-omics approach enhances the knowledge of the main molecular characteristics for each PDX model, leveraging genomic, epigenomic and transcriptomic data to support the *in vivo* drug testing on the platform.

## Results

### The ITCC-P4 PDX platform

For the ITCC-P4 consortium, a pipeline for PDX model establishment and molecular characterization was developed (**Figure 1A**; *see Methods*). Briefly, fresh patient-derived tumor samples were collected at each of the ITCC-P4 partner sites (**Supplementary Table 1**), where they were transplanted into immunodeficient mice. CNS tumor samples were injected orthotopically into the brain, while non-brain tumors were transplanted subcutaneously (flank or interscapular fat pad). Established PDX tumors were propagated through the first human-to-mouse (P0) and at least two consecutive mouse-to-mouse (P1-2) transplantation passages. Even though the molecular analyses show that most PDX models show close overlap with the matched human. Tumor fragments from established PDX models were harvested and processed for molecular characterization, together with matching patient tumors and germline material whenever available (F**igure 1A**).

**Figure 1.**
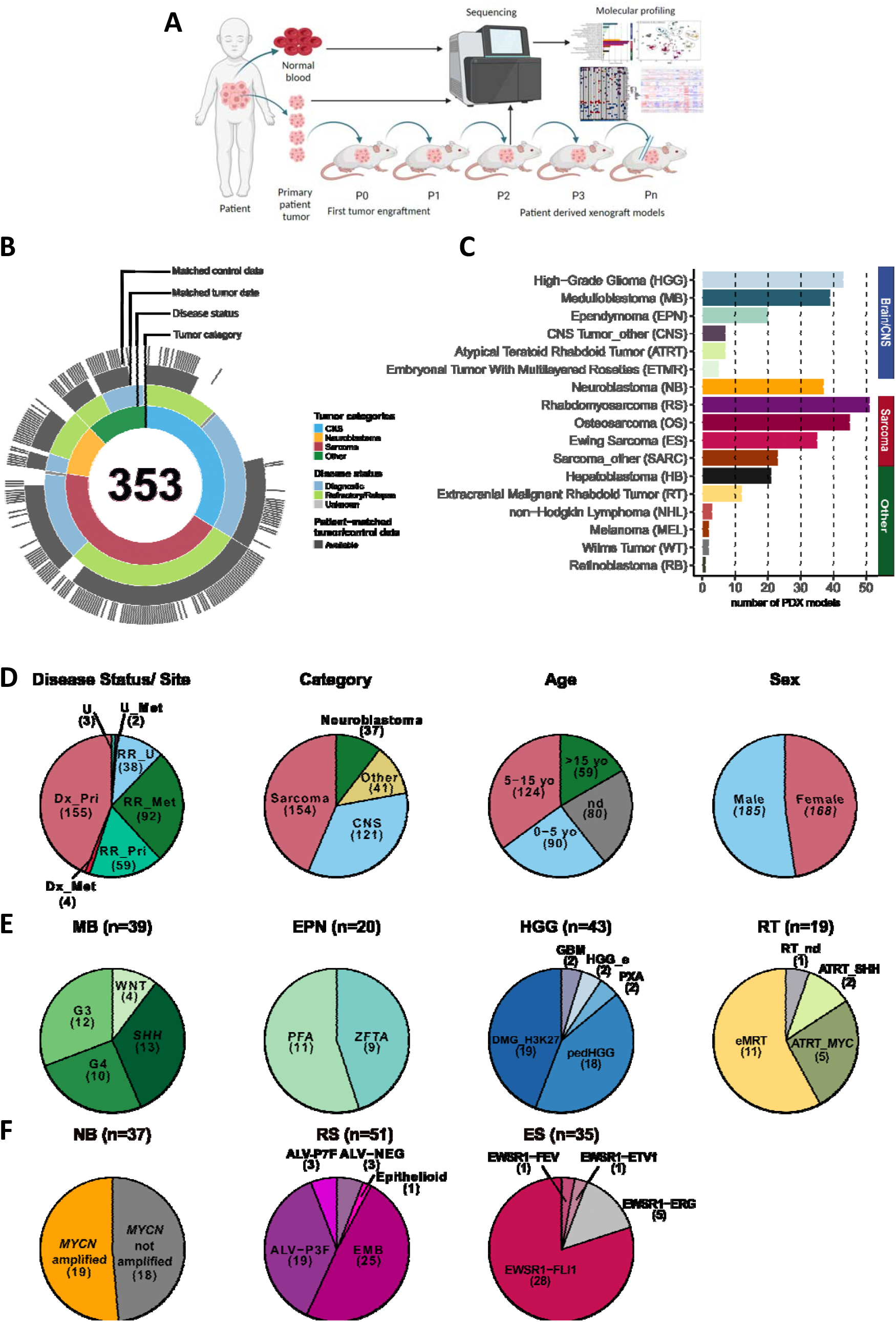
ITCC-P4 PDX models: Pipeline and Cohort Overview. A) Schematic representation of PDX establishment and molecular characterization. Tumors obtained from pediatric cancer patients were first injected into mice (first passage, P0); following two mouse-to-mouse passages (P1 and P2), the PDX models were considered established. Multiomic profiling of established PDX models and matching patient material (tumor, blood) included DNA methylation, low-coverage whole genome sequencing, high-coverage whole exome sequencing, RNA sequencing, and Affymetrix gene expression profiling. PDX tumor material harvested in the following xenopassages (P2 and higher) either supplies the ITCC-P4 PDX biobank or was used for downstream applications (drug testing, *in vitro* cultures, etc.). B) Multi-layered doughnut chart summarizing the major tumor and molecular data details of the 353 PDX models described in this study. CNS: Central Nervous System. C) Number of PDX models established for each pediatric tumor entity. D) Pie charts highlighting the main clinical features of the ITCC-P4 PDX cohort, including tumor disease status and tumor site; main cancer categories; patient age at first diagnosis; patient sex. Pri: primary; Met: metastasis; RR: refractory/relapse; U: unknown; yo: years old. E) Representation of the number of PDX models belonging to the main tumor molecular subgroups, annotated according to their epigenetic-based classification, for medulloblastoma (MB), ependymoma (EPN), high-grade glioma (HGG) and rhabdoid tumors (RT). F) Number of neuroblastoma (NB), rhabdomyosarcoma (RS) and Ewing sarcoma (ES) PDX models presenting key/typical molecular alterations (*MYCN* amplification in neuroblastoma; molecular subtpypes and FOXO*1* fusion status in rhabdomyosarcoma; EWSR*1* fusion status in Ewing sarcoma, respectively).

The PDX collection of 353 established models represents models either from diagnostic tumors (159; 45%), refractory/relapsed disease (189; 54%), or unknown disease state (5; 1%). Tumors originating from primary (214; 61%) or metastatic (98; 28%) sites were included. For 41 models (11%) this information was not available (**Figure 1B and 1D**). Additional clinical information related to the tumors used to generate PDX models, such as disease status, tumor location, patient age and sex was systematically collected (**Figure 1D and Supplementary Table 1**).

The PDX collection covers a broad range of distinct pediatric tumor types, derived from CNS (121; 34%) and non-CNS tumors (232; 66%) (**Figure 1B-D**). Amongst the CNS malignancies, we established models for medulloblastomas (MB; 39), high-grade gliomas (HGG; 43), ependymomas (EPN; 20), embryonal tumors with multilayered rosettes (ETMR; 5), atypical teratoid rhabdoid tumors (ATRT; 7) and other rare CNS cancers (CNS tumors with PLAGL1-fusion [HGNET-PLAG; 3]; choroid plexus carcinoma [CPC; 1]; CNS high-grade neuroepithelial tumor-BCOR altered [CNS-BCOR; 2]; CNS-neuroblastoma [CNS-NB; 1]). Non-CNS solid tumor models include neuroblastoma (NB; 37), sarcoma (134) (including osteosarcoma [OS; 45]; rhabdomyosarcoma [RS; 51], Ewing sarcoma [ES; 35], and additional rare sarcoma subtypes [SARC; 23]), and 41 models representing other solid pediatric malignancies (**Figure 1B-C and Supplementary Table 1**). Molecular data generated for all models (**Supplementary Figure 1B**) shows that the PDX cohort also covers the most common and representative molecular subtypes and key driver genes seen in pediatric solid cancers (**Figure 1E-F and Supplementary Table 1**).

### Tumor molecular subtyping based on DNA methylation analysis

To categorize PDX models, we first employed DNA methylation profiling. PDX models were categorized utilizing the Heidelberg CNS tumor (20) and sarcoma classifiers (21), respectively. This approach enabled us to confidently (calibrated scores >0.84, see Methods) classify 239/333 (72%) models included in the analysis (**Supplementary Table 2**).

To validate this categorization based on classifiers, as well as to classify models with low-calibrated scores and for tumor types not represented in the Heidelberg classifiers (e.g. hepatoblastoma and Wilms tumor), we clustered all methylation profiles using a t-distributed stochastic neighbor embedding (*t*-SNE) approach (**Figure 2A**). ITCC-P4 PDX (n=333) and matching patient tumor (n=193) methylation data were analyzed together with reference methylation datasets representing the same tumor types reported in our PDX cohort, supplemented with additional WHO-classified pediatric cancers (n=1169) (22–28). Most PDX models with available methylation data (333/353; 94%) clustered together with reference samples reflecting the patients’ diagnosis and/or their predicted classification (**Figure 2A**); PDX outlier samples (10/333; 3%), formed by ependymoma, hepatoblastoma, melanoma and some other rare tumor types, did not cluster with any of the representative reference samples. Primary patient tumors mostly overlapped closely with corresponding PDX within their respective “tumor type” clusters (**Figure 2A**). For the highly heterogeneous CNS tumors in our cohort, we performed sub-clustering analyses along with additional reference tumor cases reflecting the molecular subtypes included in the most recent edition of the WHO classification of Tumors of the Central Nervous System (29) (**Figure 2B, Supplementary Table 2**). The PDX models, combined with reference samples, reveal the epigenetic profiles characteristic of different molecular subtypes within various brain tumor types, including medulloblastomas (MB WNT, MB SHH, MB G3, and MB G4), ependymomas (EPN; including the two most aggressive pediatric subtypes supratentorial ZFTA-fusion positive EPN and posterior fossa PFA), high-grade gliomas (HGG) (with diffuse midline glioma, H3 K27-altered [“DMG_H3K27”], pleomorphic xanthoastrocytoma [PXA], diffuse pediatric-type high-grade glioma, RTK1 [“pedHGG_RTK1”], diffuse hemispheric glioma, H3 G34-mutant [“DHG_G34”] and high-grade glioma, MYCN [“pedHGG_MYCN”] subtypes enriched), and atypical teratoid-rhabdoid tumors (with ATRT-MYC and SHH models, but no TYR models) (**Figure 2B**).

**Figure 2:**
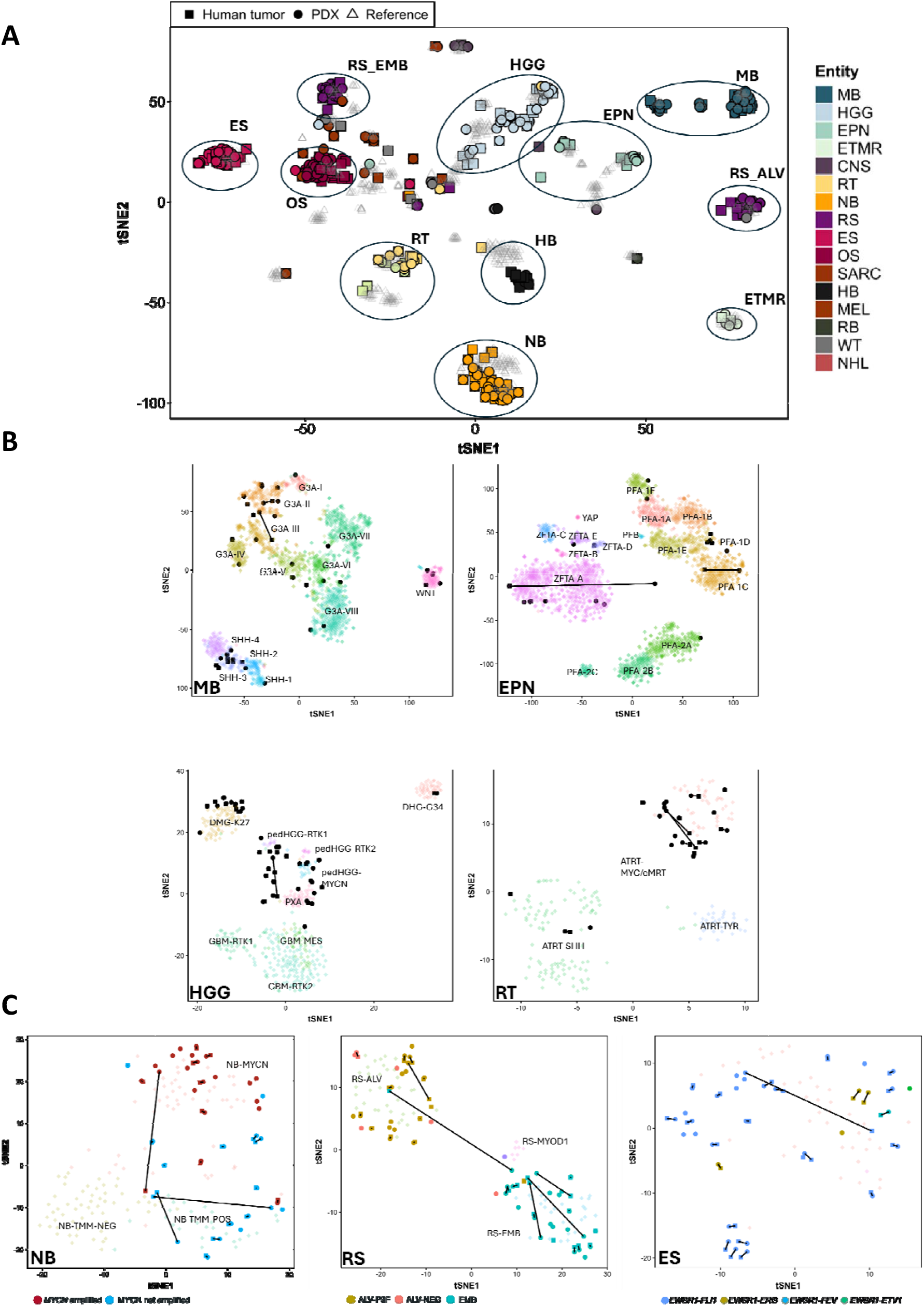
DNA methylome and transcriptome of PDX models and matching patient tumors. A) tSNE plot showing clustering analyses ITCC-P4 methylation samples (PDXs: circles; patient tumors: squares), along with external tumor references covering a broad range of pediatric tumor entities. Clusters representing the common pediatric solid malignancies are highlighted. B) tSNE panels showing subclustering analyses ITCC-P4 methylation data grouped with consensus references for the main pediatric brain/CNS (medulloblastoma, ependymoma, high-grade glioma) and rhabdoid tumor entities with relative molecular subtypes. Connecting lines are used to link PDX models (black dots) with their matching human tumor (black squares) data. C) Subclustering analyses of neuroblastoma (left panel), rhabdomyosarcoma (middle panel) and Ewing sarcoma (right panel) methylation datasets, including ITCC-P4 and reference samples. Matching PDX and tumor data are connected. Key molecular status, like *MYCN* amplification and detected *EWSR1* fusion events are highlighted in the respective plots.

For neuroblastoma (NB), PDX models formed two separate clusters according to their *MYCN* amplification status (**Figure 2C, left**). Rhabdomyosarcoma (RS) models showed a clear separation between alveolar and embryonal subtypes, both well-represented in our PDX cohort (**Figure 2C, middle**). Ewing sarcoma (ES) PDX models co-clustered with ES reference tumors (**Figure 2C, right**). The methylome profiles of the PDX samples demonstrated concordance with those of the corresponding tumor samples, frequently clustering as the nearest neighbors in the clustering analysis. Exceptions of higher distance within the clusters, suggesting dissimilar profiles, were observed in one EPN, two NB, one RS and one ES PDX-tumor pairs (**Figure 2B-C**). Models with unusual molecular features and the remarked difference in tumor cell purity between patient and PDX samples (**Supplementary Table 2**), as a result of remaining mixed murine and human reads of the PDXs, could lead to the divergent clustering behaviors observed in these latter pairs.

To infer epigenomic correlations between the PDX models and corresponding patient tumors, we calculated a similarity index between the available PDX-tumor pairs (n=184). Manhattan distance calculated for each PDX-tumor pair demonstrated an overall high epigenetic similarity over the SNP sites (157/184) of comparisons with distance <0.1) and, therefore, the capacity of the PDX models to recapitulate the epigenetic milieu observed in human tumors (**Supplementary Figure S2A**).

### Transcriptomic analysis of tumors and PDX models

Next, we analyzed PDX and tumor transcriptome data, focusing on grouping samples based on their similarity in gene expression patterns. Gene-based (top 1000 most variable genes across the cohort) hierarchical clustering of PDX transcriptome data showed that our models tended to aggregate based on their tumor type; in particular, co-clustering of PDX models was driven by the expression levels of key genes found differentially deregulated in pediatric cancers, such as e.g. *MYCN, ALK* (upregulated in NB), *MYOD1* (expressed in RS) or *PAX3* (expressed in ES) (**Supplementary Figure S2B and Supplementary Table 3**). A comparison of patient and PDX tumor samples, based on differentially expressed genes, revealed a consistent preservation of the tumor-core signature from the original human tumors to the PDX models. In comparison with patient transcriptomic data, PDXs were generally characterized by downregulation of genes related to immune and stromal functions (**Supplementary Table 3**). These results corresponded to predicted tumor purity scores calculated on PDX and tumor samples based on omics data (**Supplementary Table 2 and 5**), in line with previous reports (8,10,30).

### The mutational landscape of the ITCC-P4 PDX models

Next, we investigated the mutational landscape of the PDX models by performing comprehensive genomic analysis. This included analysis of somatic single-nucleotide variants (SNVs), small insertions/deletions (indels), structural variations (SV), copy number variations (CNVs) and gene fusion events from the PDX genome (**Figure 3 and Supplementary Figure S4E**), in combination with the matching human tumors (**Supplementary Figure S3, Supplementary Table 4**). Germline mutation calling was performed whenever matched control material was available (195/353; 55%). The full repertoire of genome-wide human tumor and PDX samples is provided here: https://r2platform.com/itcc-p4.

**Figure 3:**
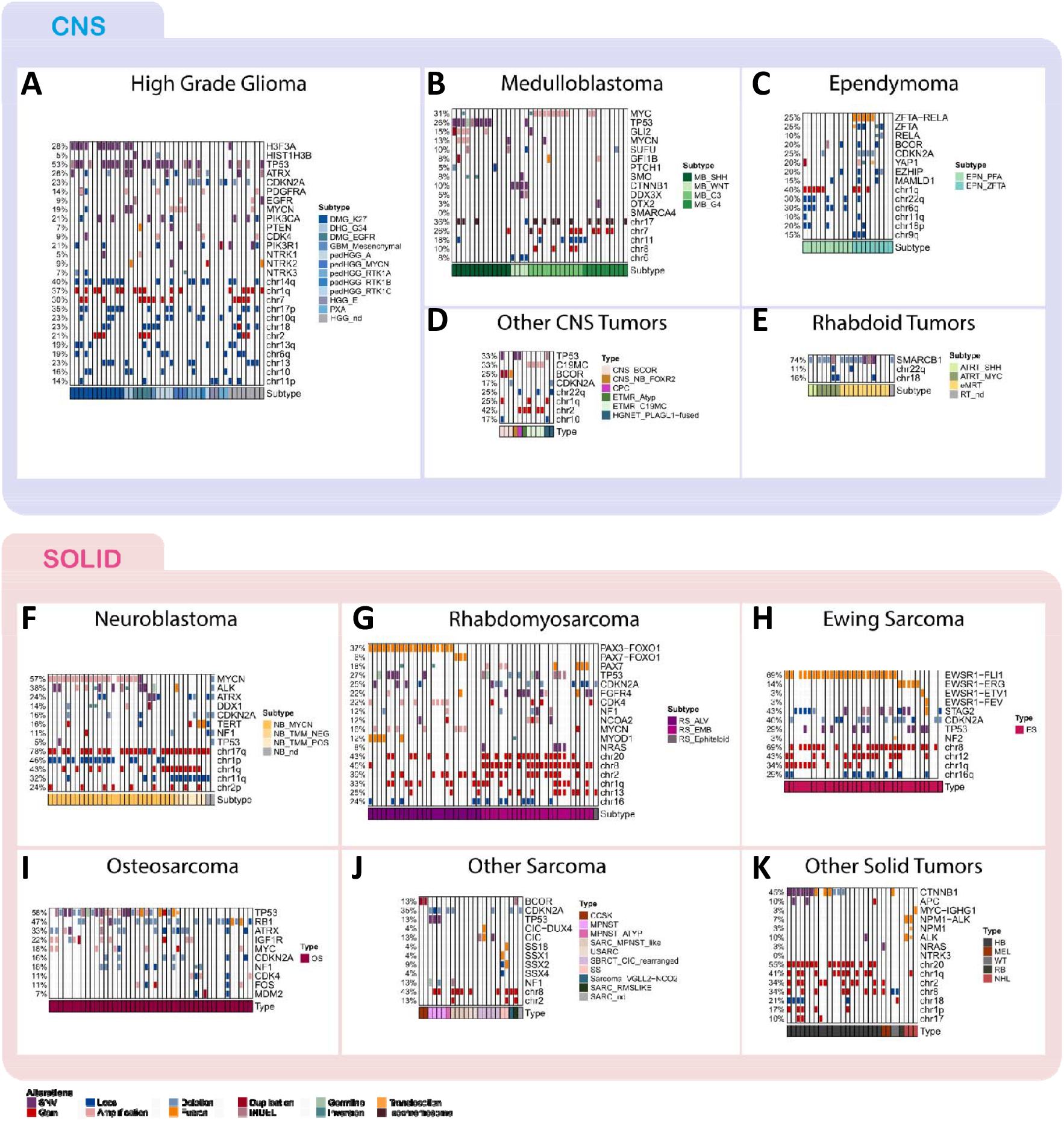
Oncoprints depicting multiple genomic aberrations in the ITCC-P4 PDX cohort. Oncoprints reporting the mutational landscape of the PDX models established within the ITCC-P4 project. Each panel shows a comprehensive view of the key genetic alterations, including somatic and germline mutations (SNVs and INDELs), structural variants (SVs) and copy number variations (CNVs), for PDX models belonging to distinct tumor types: High-grade glioma (A); Medulloblastoma (B); Ependymoma (C); Other CNS tumors (D); Rhabdoid tumors (E); Neuroblastoma (F); Rhabdomyosarcoma (G); Ewing sarcoma (H); Osteosarcoma (I); other sarcomas (J); other solid tumors (K). Tumor subtypes for each PDX model, if known and defined based on the methylation analyses, were reported.

We calculated the total number of somatic mutations within the exome present in PDX and tumor samples to define the tumor mutational burden (TMB). We observed a high concordance of the TMB between PDXs and corresponding tumors (**Supplementary Figure S4A-C**); a slightly higher TMB was detected in refractory cases compared to relapse and diagnostic states, with a greater difference seen between PDX and Tumor cases of diagnostic samples. Metastatic tumor samples showed an overall higher TMB compared to metastatic cases (**Supplementary Figure S4B-C**). Notably, HGG, EPN, RT, SARC, OS, HB, WT entities demonstrated a pronounced disparity in TMB between the tumor and paired PDX, with the latter showing a higher TMB (Sup**plementary Figure S4D**).

To prioritize functionally and clinically relevant SNVs, we then focused on a curated set of known tumor type-specific driver genes, selected based on their involvement in tumorigenesis and their potential as therapeutic targets (31,32). The full report of annotated variants (called with the hg19 assembly) per each model are available on our public platform ITCC-P4 Data scope in R2 (https://r2platform.com/itcc-p4) upon request.

#### CNS tumors and ATRT/extracranial rhabdoid tumor PDX models

PDX models generated from pediatric CNS malignancies harbored recurrent mutations that reflected the predicted molecular subtypes based on DNA methylome profiling (**Figure 3A-E**).

Pediatric HGG is a heterogeneous family of tumors, represented in our PDX cohort (n=43; Figure 3A) by models of DMG_H3K27 (19/43, including n=4 *EGFR*-mutant cases, “DMG_EGFR” subtype), pediatric high-grade gliomas (“pedHGG”; 18/43, including the “pedHGG_A” [n=4], “pedHGG_MYCN”[n=3], “DHG-G34” [2/43], “pedHGG_RTK1” [n=3], “pedHGG_RTK2” [n=1] and”pedHGG_RTK3” [n=1]; n=4 with unknown subtype), PXA (2/43), glioblastoma (“GBM”; 2/43, including “GBM_MES” [n=1] subtype and n=1 with unknown subtype) and adult-type diffuse high-grade glioma, subtype E (“HGG_E”; 2/43). The majority (30/43) were established from primary diseases. The most frequently mutated gene in this group was *TP53* (23/43), followed by *H3F3A* (12/43 mostly in DMG PDXs), *ATRX* (11/43), *CDKN2A* (10/43) and *MYCN* (8/43). Moreover, recurrent mutations in effectors of signaling pathways, such as *BRAF, NRAS, PIK3CA, PTEN, EGFR* and *MYB*, and structural variations affecting *NTRK1*/2/3 were also identified in these models.

The 39 MB PDX models (**Figure 3B**), representing all four main molecular groups (4 WNT, 13 SHH, 12 G3, and 10 G4), all displayed subgroup-specific aberrations. The majority (30/39) of MB models were established from primary lesions. The other nine were from relapsed disease (5 SHH, 1 G3, 2 G4) or unknown (1 G3). *MYC* amplification predominantly occurred in G3 models (8/12 total G3), while SHH models showed recurrent *TP53* mutations (9/13), including one case with a confirmed *TP53* germline alteration. Other SHH-specific gene variants included *GLI2* (4/13) and *MYCN* (3/13) amplifications.

The 20 EPN PDX models represented only two of the most aggressive molecular subtypes in EPN disease: posterior fossa type A (PFA EPN; 11) and supratentorial ZFTA-fused EPN (ZFTA; 19). In 7/9 ZFTA models (so classified based on the methylation analyses), the ZFTA locus was shown to be affected by either fusion events (ZFTA-RELA, 5), inversion (1), or focal loss (1), suggesting a predominant ZFTA-fusion driven status in this subtype, although we could not formally prove this in all models. Interestingly, most of the established models (17/20) were established from refractory/relapsed disease. High-risk features were also clearly overrepresented in these models: 1q gain in 8/20 EPN models, 6q loss exclusively found in PFA models (6/11), and CDKN2A loss in 5/20 models (**Figure 3C**).

Our PDX cohort further included n=5 ETMR PDX models (4/5 presenting the typical C19MC amplification) (**Figure 3D**) and seven CNS embryonal tumors that were classified, based on their methylation profiles (**Supplementary Figure 2**), as BCOR-altered tumors (n=3; the typical *BCOR* in tandem duplication observed in 2/3 models, while in 1/3 showed a *BCOR* fusion), HGNET PLAG1-fused (n=2), CNS-neuroblastoma (n=1) and CPC (n=1). In this small cohort, genes like *TP53* and *CDKN2A* were frequently observed as mutated, as well as chr1q and 2 gains (**Figure 3D**).

PDX models of CNS atypical teratoid/rhabdoid tumors (ATRTs, 6/7 of diagnostic disease, 1/7 from refractory tumor; all from primary lesions) and extracranial malignant rhabdoid tumors (eMRTs, n=12, with 11 from primary sites and 1 from metastases), were mainly characterized by *SMARCB1* alterations (SNVs n=2/19; INDELs n=2/19; copy loss n=10/19; undetected in 5/19) and exhibited, as expected, very few other genetic alterations (**Figure 3E**).

#### Neuroblastoma PDX models

Among the 37 NB models (10 from diagnostic primary tumors, 4 from relapse primary site, 3 from refractory primary site, 1 from refractory metastasis, 13 from relapse metastasis, 6 from relapse tumor with undefined site), *MYCN* alterations were present in the majority of the PDXs (21/37), most frequently as focal gene amplification (19/21) (**Figure 3F**). Additional neuroblastoma-associated driver mutations affecting *ALK, ATRX, CDKN2A* and *DDX1* were also found (with 14, 9, 6 and 5/37 cases, respectively). *NF1* (4/37), *BRAF* (4/37), *TP53* (2/37) mutations were also observed. The most frequent CNVs observed in NB models were chr 17q gain (29/37) and 1p loss (17/37). Additionally, we also observed CNVs linked to aggressive disease and poor outcome (33,34) such as 1q loss (16/37) and 11q loss (12/37).

#### Soft-tissue sarcoma/*Bone sarcoma PDX models*

Among the 51 RS PDX models, both alveolar (25; 4 diagnostic/primary, 2 relapse/primary; 14 relapse/metastasis; 3 refractory/primary; 1 refractory/metastasis; 1 relapse/undefined site) and embryonal (25; 10 diagnostic/primary, 5 relapse/primary, 4 relapse/metastasis, 2 refractory/primary, 1 refractory/metastasis, 2 relapse/undefined site and 1 unknown) subtypes were represented. One model was annotated as epithelioid based on the pathological findings and presented a distinct molecular status. Genetically, a clear distinction was observed between alveolar RS models with *PAX3-FOXO1* fusions (“ALV-PF”; 19/25), which is the key alteration in the alveolar subtype, while 3 models displayed a *PAX7-FOXO1* fusion. *CDK4* amplification or gain was also only identified in alveolar RS models (11/25). Embryonal models harbored more frequent alterations in *TP53* (9/25), *NRAS* (4/25) 19, *NF1* (4/25), *BCOR* (8/25), *FGFR4* (9/25), *NCOA2* (4/25) and *CDKN2A* (8/25) (**Figure 3G**).

Analysis of the 35 ES PDX models (13 diagnostic/primary, 1 diagnostic/metastasis, 4 relapse/primary, 8 relapse/metastasis; 4 refractory/primary; 2 refractory/metastatic, 2 relapse/undefined site; 1 unknown; **Figure 3H**) revealed, as expected, *EWSR1::FLI1* fusion as the most recurrent aberration (28/35). Other known *EWSR1::ETS* fusion genes were also observed, such as *EWSR1-ERG* (5/35), *EWSR1-FEV* (1/35) and *EWSR1-ETV1* (1/35). Additional mutations identified included recurrent SNVs associated with poor outcomes in *STAG2* (15/35) and *TP53* (10/35), co-occurring in five models (35). *CDKN2A* loss or deletion was identified in 14/35 ES PDXs. 22/35 ES models exhibited the typical whole chr 8 gain, and other known CNVs for this tumor entity (35), such as chr1q gain (12/35), chr12 gain (15/35), chr16q loss (10/35), were also frequently observed in our cohort.

OS models (45; 13 diagnostic/primary; 2 diagnostic/metastasis; 1 relapse/primary; 16 relapse/metastasis; 3 refractory/primary; 7 refractory/metastasis; 3 relapse/unknown) reflected the high genetic complexity observed in human tumors with the detection of many somatic variants (**Figure 3I**). Mutations in *TP53* (26/45) and *RB1* (21/45) indeed occurred frequently in the OS PDX cohort, followed by *ATRX* loss/deletion (12/45), *MYC* gain/amplification (7/45), *IGF1R* SVs/CNVs (10/45), *NF1* mutations (8/45) and *CDKN2A* loss/deletion (7/45). As expected, the CNV landscape of these models was highly complex, with each case displaying a unique and complex pattern of CNV alterations (**Supplementary Figure S4E**).

PDX models representing other rare sarcoma types, such as malignant peripheral nerve sheath tumors (MPNST; 1 diagnostic/primary, 1 refractory/primary, 2 relapse/metastasis), clear cell sarcoma of the kidney (CCSK; 1 relapse/metastasis, 1 relapse/undefined site), synovial sarcoma (SS; 1 relapse/primary, 1 relapse/metastasis), CIC-rearranged soft tissue round cell sarcoma (SBRCT_CIC_rearranged; 2 diagnostic/primary, 1 refractory/primary, 1 relapse/metastasis, 1 relapse/unknown) and undifferentiated sarcoma (1 relapse/primary, 1 relapse/metastasis) displayed recurrent alterations such as *CDKN2A* loss, *BCOR* internal tandem duplication (in both CCSK models) and *TP53* mutations. In SS models, we detected fusions of *SSX* and SS family member genes *SSX1, SSX2, SSX4* and *SSX18*, while structural variants or fusion events involving *CIC* (in particular *CIC::DUX4*) were exclusively observed in SBRCT_CIC_rearranged models (**Figure 3J**).

#### Other PDX models

The remaining PDX models represent a heterogeneous group of rare pediatric tumors of which the largest group is the hepatoblastoma (HB, 21) models presenting recurrent beta-catenin gene (*CTNNB1*) mutations (12/21), as well as typical chr1q gain, whole chr20 gain and chr18 loss. Other rare models include two models for pediatric melanoma (MEL), one of which with a *NRAS* mutation. 1/2 of Wilms tumor (WT) PDXs exhibited a *CTNNB1* fusion, while three non-Hodgkin lymphoma (NHL) models were mostly characterized by fusions involving *ALK* or *MYC* (**Figure 3K**).

### Assessing PDX model fidelity compared to matching patient tumors

Next, we compared molecular data from matched human tumor samples and PDX to evaluate fidelity and identify genomic landscape disparities (**Supplementary Figure S3, Supplementary Table 4**).

When comparing the genomic data of patient tumors and paired PDXs (number of pairs: 136; PDX models without matching patient tumor and/or germline WES data could not be included in this analysis), we first considered that tumor samples obtained from patients and xenografts might have different tumor cell content. For this reason, we evaluated this aspect using distinct metrics: Tumor cell fraction (TCF) scores from CNVKit. ploidy (both derived from WES data analysis) and Estimation of STromal and Immune cells in MAlignant Tumor tissues using Expression data scores (ESTIMATE score derived from RNAseq data). Based on the TCF scores, we observed that, while patient tumor samples exhibited high variability, most PDX models (99/136, 73%) showed the highest (1) TCF score, confirming the expected difference in tumor cell content (**Supplementary Figure S5A and Supplementary Table 5**).

Next, we compared the variant allele frequencies (VAF) of driver gene SNVs between human tumors and matched PDXs, following a correction of the VAFs for each sample (tumor or PDX) based on their TCF. Overall, human tumor-PDX pairs showed a high concordance of TCF-corrected VAF values for shared somatic variants (Pearson’s correlation >=0.2 in 95 PDX-tumor pairs out of a total of 136 comparisons) (**Supplementary Table 5**).

Changes in corrected SNV VAFs from patient tumors to xenografts were defined using the delta VAF parameter (*see Methods*). First, we focused on the most frequent somatic mutation events (*ALK, ATRX, BRAF, BCOR, CDKN2A, FGFR4, H3F3A, NTRK3, NOTCH3, PIK3CA, PTPNN11, RB1, TP53*); 65/136 tumor/PDX pairs presented a mutation for one or more of these selected genes (total 142 gene comparisons). The delta VAF scores for these variants indicated that 70% of the cases (100/142) shared somatic mutations presented stable VAFs between human tumor and PDX (delta VAF scores ranging from -0.2 and 0.2). In some cases, we could detect higher VAFs in human tumors (negative delta VAF <-0.2; 8/142) or PDXs (positive delta VAFs >0.2, 34/142). As an example, *TP53* SNVs were the most frequent somatic mutations with 36/65 PDXs showing a comparable VAF to the respective tumors. However, in some instances, we could detect either a tumor-(2/36) or PDX-(10/36) VAF-score enrichment for TP53 and other key drivers (Fig**ures 4 A-D**; Supplementary Figure S5B): HGG PDX models frequently presented higher *TP53, PIK3CA* and *ATRX* VAFs compared to their matching tumors, while *BRAF* mutations in HGGs were seen exclusively in the human tumors and absent in the PDXs, suggesting a selection of a non-BRAF -mutated clone upon PDX establishment (**Figure 4A**). NB and RS PDX models presented SNVs with VAFs strongly concordant with their matching tumors (**Figure 4B-C**), as in the NB model NB0025 (*ALK, MYCN* and *BRCA2* mutated) and the RS model RS0189 (TP53 and ALK mutated) (**Figure 4E**). A few exceptions concerned with key mutations observed exclusively in patient tumors and not found in the PDXs, such as *ALK* and *ATRX* in two NB models (**Figure 4B**), or *FGFR4* and *CDKN2A* in two and one RS models, respectively (**Figure 4C and 4E**). An important increase in VAF was also found in OS PDX models exhibiting *TP53* and *RB1 SNVs*, while tumor models showed higher VAF than PDX with BRAF and *ATRX SNVs* (**Figure 4D**). Finally, we observed rare PDX models that presented a remarkable divergence from the matching human tumor. For example, in the HGG PDX HG0354 and its matched tumor most of the selected SNVs showed a mutually exclusive pattern, including the mutation affecting *TP53*, observed only in the PDX sample, with only a few other mutations shared between the two conditions (**Figure 4E**).

**Figure 4:**
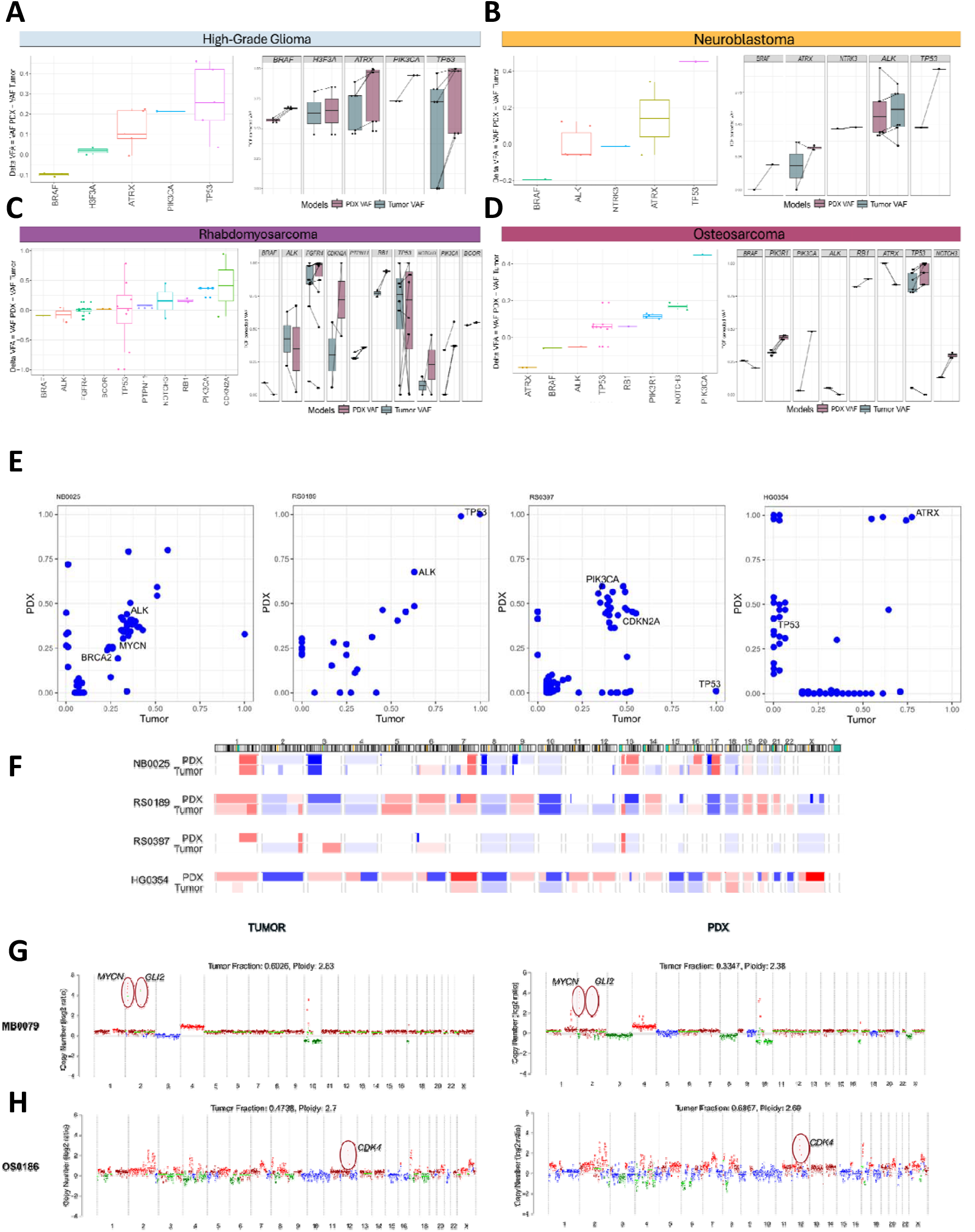
Comparison of molecular alterations between patient tumors and matched PDXs. A) On the left panel: delta VAF scores (on the y-axis; calculated by subtracting the TCF corrected VAF of matching patient tumor from the TCF corrected VAF of the PDX model) calculated for key driver genes (*BRAF, H3F3A, ATRX, PIK3CA, TP53*) in high-grade glioma tumor and PDXs. Positive deltaVAF scores indicate a higher frequency of the given mutation in the PDXs compared to their matched human tumors, while negative values define a higher mutation frequency in the original tumor. Highly concordant PDX-tumor pairs show deltaVAF scores close to zero. On the right panel: TCF corrected tumor and PDX VAFs (y-axis) associated with the somatic SNVs affecting the most frequently mutated driver genes in high-grade glioma. Black lines connect matching pairs. B) Delta VAF (left) and TCF corrected VAF (right) plots showing the comparison of the mutation frequencies for key mutated drivers (*BRAF, ALK, NTRK3, ATRX, TP53*) in neuroblastoma PDX and tumor samples. C) Delta VAF (left) and TCF corrected VAF (right) plots showing the comparison of the mutation frequencies for key mutated drivers (*BRAF, ALK, FGFR4, BCOR, TP53, PTPN11, NOTCH3, RB1, PIK3CA, CDKN2A*) in rhabdomyosarcoma PDX and tumor samples. D) Delta VAF (left) and TCF corrected VAF (right) plots showing the comparison of the mutation frequencies for key mutated drivers (*ATRX, BFRA, ALK, TP53, RB1, PIK3R1, NOTCH3, PIK3CA*) in osteosarcoma PDX and tumor samples. E) Scatter plots showing the variant allele frequencies (VAFs) of somatic SNVs shared between the PDX models NB0025, RS0189, RS0397, HG0354, and their respective patient tumors. Key mutated genes are highlighted. F) Heatmap of the whole-genome CNV landscape observed in the PDX models NB0025, RS0189, RS0397, HG0354 and their respective patient tumors. Gains of chromosome regions are shown in red, while region losses are represented by blue segments. G) Genome-wide plots (generated by ichorCNA) showing the copy number ratio (log2) and focal amplification events of *MYC* and *GLI2* in MB0079 PDX model (right) and tumor (left). Tumor fraction and ploidy are reported for both samples; H) Genome CNV plots of OS0186 PDX (right) and matching tumor (left). CKD4 amplification (red circles) is observed in both conditions;

Next, we compared large-scale CNVs and structural variants (SVs) in human tumor/PDX pairs (**Supplementary Figure S3**); PDX models exhibited, again, an overall strong concordance with their respective tumors based on their CNV profiles. However, PDX models such as RS0397 and HG0354 presented divergent CNV profiles as compared to patient tumors, in line with the different mutational profiles (**Figure 4E-F**).

Overall, the PDXs included in this study exhibited good molecular concordance with their respective human tumors in terms of variant allele frequencies, while mutually exclusive alterations were sporadically observed.

### Modeling Tumor Progression: “serial” PDX models

An important feature of the ITCC-P4 PDX resource is the availability of serial PDX models, providing insights into the underlying mechanisms driving tumor progression. In our cohort, serial PDX models were successfully established from 40 tumors collected from 19 patients (2-4 models generated per patient). This model series included: primary/diagnostic and post-treatment progressive tumor-derived PDXs (seven patients), primary/initial and relapse (or multiple relapses) PDXs (five patients), a refractory/recurrence PDX pair and PDXs models established from different patient-derived metastases (three patients) (Figure 5A). Multi-omics molecular characterization of these models revealed distinct molecular features mirroring the specific events in each patient’s tumor. A detailed overview of the key molecular alterations detected in the serial PDX model is included in **Figure 3** and **Supplementary Figure S3**.

**Figure 5:**
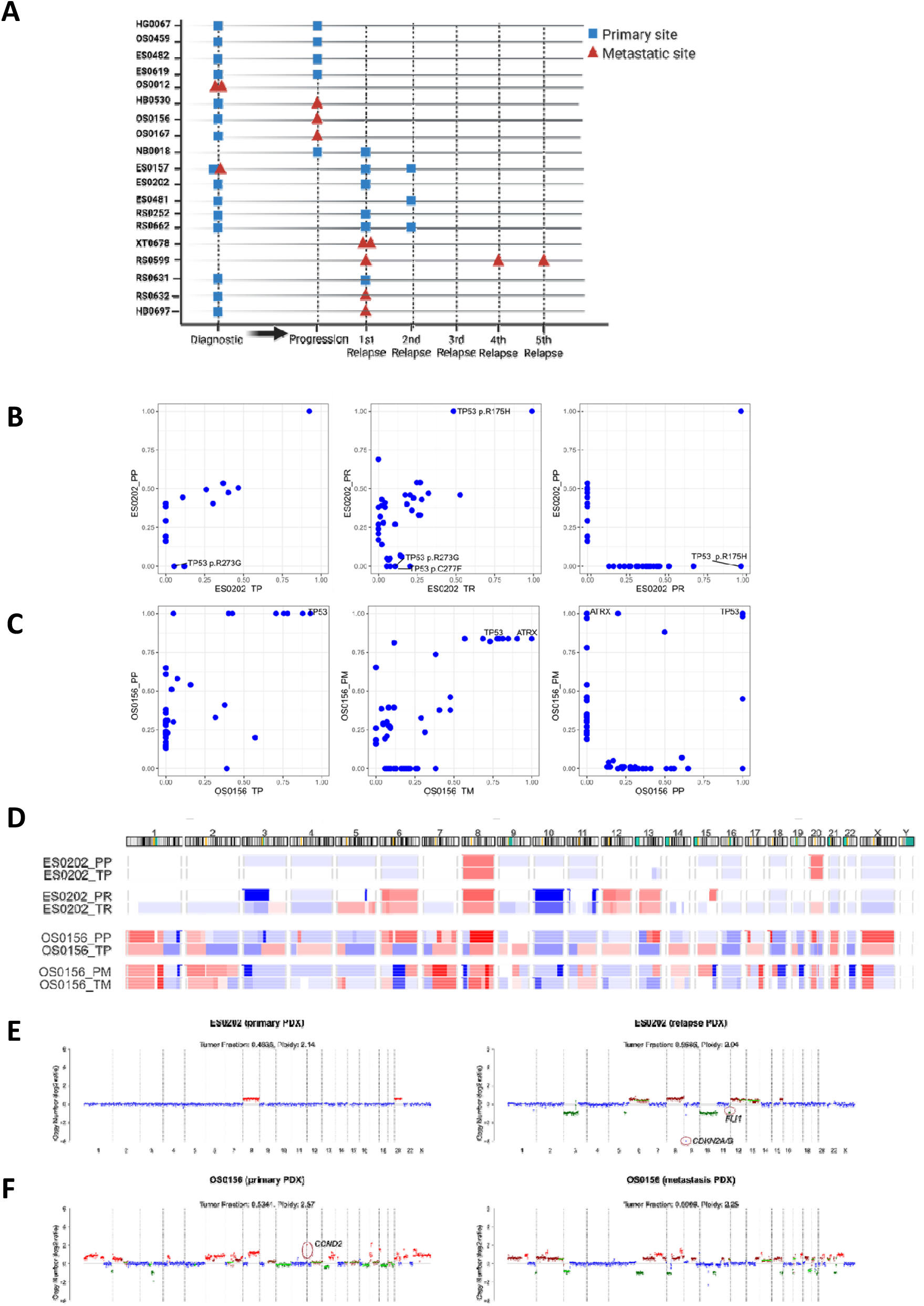
Comparative analysis of the “serial” PDX models reveals tumor event-specific molecular characteristics. A) Schematic overview of the subset (n=40) of serial PDX models established from multiple patient tumor events over the course of the disease. The tumor site from which multiple PDX were generated was highlighted; B) VAF plots showing the correlation of mutation frequencies for the following combinations of the Ewing sarcoma models ES0202: patient primary tumor (PT) vs. primary PDX (PP; left); patient relapse original tumor (TR) vs. relapse PDX (PR; center); relapse PDX vs. primary PDX (right). TP53 gene mutations are highlighted; C) VAF plots of models of the osteosarcoma models OS0156 under the following combinations: patient primary tumor (PT) vs. primary PDX (PP; left); patient metastatic tumor (TM) vs. metastasis PDX (PM; center); metastasis PDX vs. primary PDX (right). TP53 and ATRX gene mutations are highlighted; D) CNV heatmap showing chromosome gains and/or losses for PDX models ES0202 (primary and relapse), OS0156 (primary and metastasis) and relative matched patient samples; E) CNV plots of ES0202 (primary and relapse) PDX models. Key focal events are highlighted in red circles; F) CNV plots of OS0156 (primary and metastasis) PDX models. Key focal events are highlighted in red circles.

Overall, serial PDX models exhibited closely aligned molecular characteristics, evident in their clustering patterns derived from their DNA methylome profiles; however, two cases (ES0202, OS0156) showed distinct clustering behavior (**Supplementary Figure S6 A-E**), suggesting different tumor evolution patterns.

Divergent genomic landscapes were observed in a few serial models. The serial models ES0202 were generated from the primary and recurrent tumors of an Ewing sarcoma patient (**Figure 5A**). Molecular characterization performed on the primary (PP) and recurrent (PR) PDX models, along with the respective patient tumor samples (TP and TR) showed a *TP53* mutation R175H, frequently observed in different tumor types and actively promoting tumorigenic features and drug resistance, that was only detected in post-treatment relapse-derived samples (TR and PR), while absent in the primary tumor and PDX (**Figure 5B**). A similar acquisition of *TP53* SNV in models generated at a later disease timepoint was observed in other ES models (ES0482, ES0t619) (**Supplementary Table 4**).

In another set of serial PDXs (OS0156) generated from a diagnostic primary OS and a metastasis detected upon progression following treatment (**Figure 5A**), we observed shared *TP53* SNV across all the tumor and PDX samples, while an *ATRX* somatic mutation could be only detected in patient metastatic tumor (TM) and the corresponding metastasis PDX (PM) (**Figure 5C**).

With regards to CNVs, serial PDX models generated from later disease stages frequently revealed novel CNVs, such as gain of chr6 and loss of chr10 in ES0202_PR or the loss of chr6q16-23, frequently observed in OS, in OS0167_PM (**Figure 5D**). Moreover, *CDKN2A/B* deletion and *FLI1* loss were exclusively detected in ES0202_PR (**Figure 5E**), while *CCND2* gain, shown in OS0156_PP, could not be seen in the matching metastasis-derived PDX model (**Figure 5F**).

In only one instance was a driver event, a PIK3CA mutation, seen in a primary PDX (RS0632_PP) but not in the corresponding relapse model (RS0632_PR) (**Supplementary Table 4**).

Altogether, serial models revealed a persistence of driver events across the disease course, with a frequent emergence of new driver alterations such as TP53 upon relapse.

### *In vivo* drug sensitivity using the R2 Data scope: Implications for targeted therapies

The ITCC-P4 platform, with its large repertoire of PDX models of high-risk pediatric cancers and multiple models per tumor type, allows *in vivo* drug testing in single mouse clinical trials. Drug response in each model can subsequently be directly correlated with all molecular data to identify biomarkers associated with response or resistance. As proof-of-concept, we evaluated the efficacy of standard of care (SOC) drugs and the ALK inhibitor lorlatinib in 18 NB and 22 RS models.

NB models with *MYCN* amplification are more likely to exhibit favorable responses to SOC drug topotecan compared to those without *MYCN* amplification. Indeed, eight NB models (seven with *MYCN* amplification) responded to topotecan (min. RTV < 1%). Another seven models showed a partial response (min. RTV < 50% but > 1%), with one model having a *MYCN* amplification and another displaying a *MYCN* mutation. Of the three models that were unresponsive (min. RTV > 50%), two lacked *MYCN* amplifications/mutations (Figure 6). Conversely, SOC drug etoposide, a topoisomerase II inhibitor, exhibited greater efficacy in models lacking MYCN amplifications (**Figure 6**). Lorlatinib was most effective in NB0457_PR and NB0277_PR (min. RTV <=1%), both exhibiting ALK amplification and elevated levels of *ALK* expression, which was absent in the 14 models with low lorlatinib sensitivity (min RTV> 50%), suggesting *ALK* amplification/expression as a predictive biomarker for lorlatinib sensitivity in neuroblastoma (36). Interestingly, four NB models with hotspot *ALK* F1174L and R1275Q mutations were not sensitive to lorlatinib.

**Figure 6:**
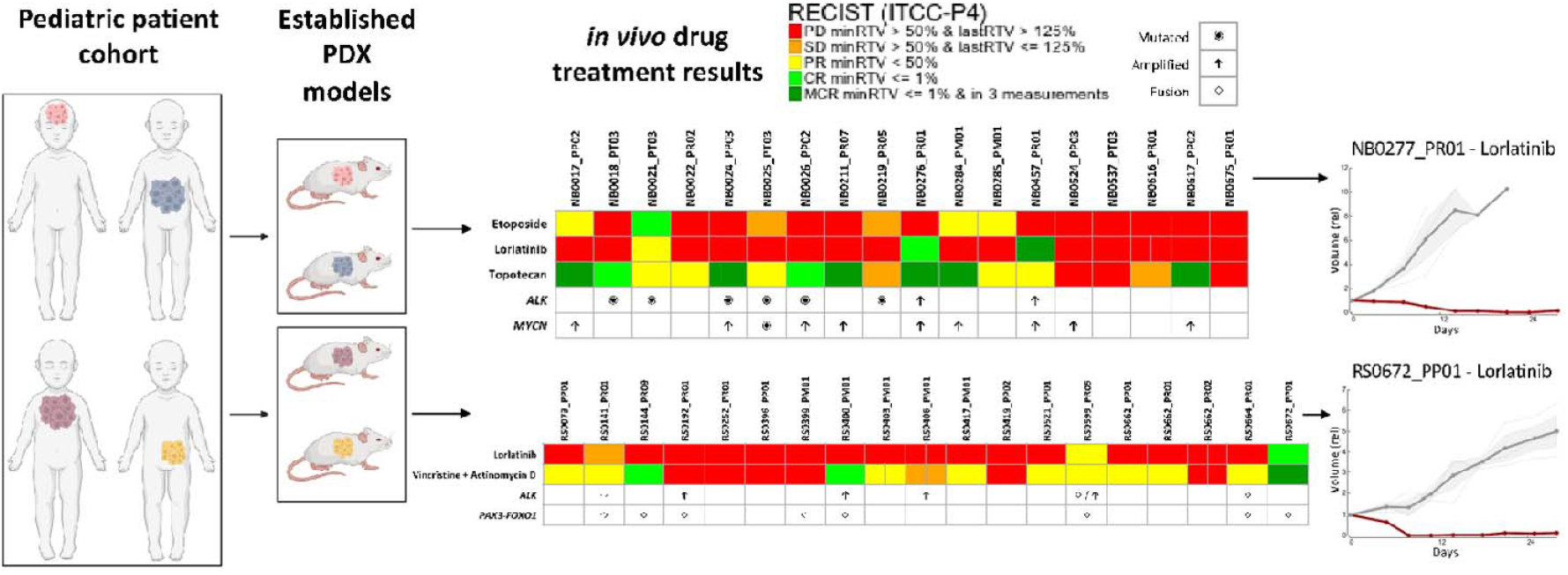
*in vivo* drug testing based on molecular characterization on R2. The ITCC-P4 PDX cohort on the R2 datascope acts as a powerful resource to allow extraction of *in vivo* drug testing data for various Standard-of-Care drugs (SOC) and ALK inhibitor Lorlatinib across NB and RS models with *in vivo* data. Lorlatinib and Topotecan across Neuroblastoma showing responders and non-responder PDX models. NB0277_PR shows a decrease in tumor growth with Lorlatinib treatment. Treatment of Lorlatinib on the Rhabdomyosarcoma samples showed effective decrease in tumor size in sample RS0672_PP.

In RS models treated with the SOC drugs vincristine and actinomycin D, 3/22 models exhibited high sensitivity (minimum RTV < 1%), and 12/22 showed a partial response (minimum RTV < 50%). Among these 15 responsive models, eight (53%) harbored the *PAX3-FOXO1* fusion gene including one model with *PAX7-FOXO1* fusion. Among the seven models with low sensitivity, only two harbored this gene fusion. Lorlatinib demonstrated high efficacy (RECIST score ≤1%) in two models (RS0599_PR05 and RS0141_PR01) harboring *ALK* fusions, suggesting that *ALK* fusions are a biomarker for lorlatinib response in RS. Interestingly, RS0672_PP also showed a very high sensitivity (minimum RTV ≤ 1%) despite the absence of any identified ALK aberrations (**Figure 6**). Further study of the correlation between molecular characteristics and sensitivity to lorlatinib in NB and RS models will be possible based on pending drug sensitivity profiles.

These results underscore the importance of understanding the tumor’s genetic profile to tailor treatment and explore additional targeted therapies that may be more effective in the context of specific genetic alterations.

### The ITCC-P4 data portal

Extensive molecular characterization data from the ITCC-P4 PDX cohort is available to the global scientific community via the R2 ITCC-P4 PDX Data scope portal (https://r2platform.com/itcc-p4/) upon access request. This portal allows researchers in an accessible user-friendly way the selection of specific models by entity, oncogenic drivers, or other biomarkers for proof-of-concept *in vivo* studies in PDX models available to the global scientific community. This includes the entire curated ITCC-P4 PDX multi-omics dataset of the PDX models described in this study as well as additional models that were generated and characterized within the ITCC-P4 project. This resource enhances the translational potential of tailored *in vivo* studies, potentially accelerating the discovery and validation of novel therapeutic targets or treatment modalities and biomarkers (**Figure 7**).

**Figure 7:**
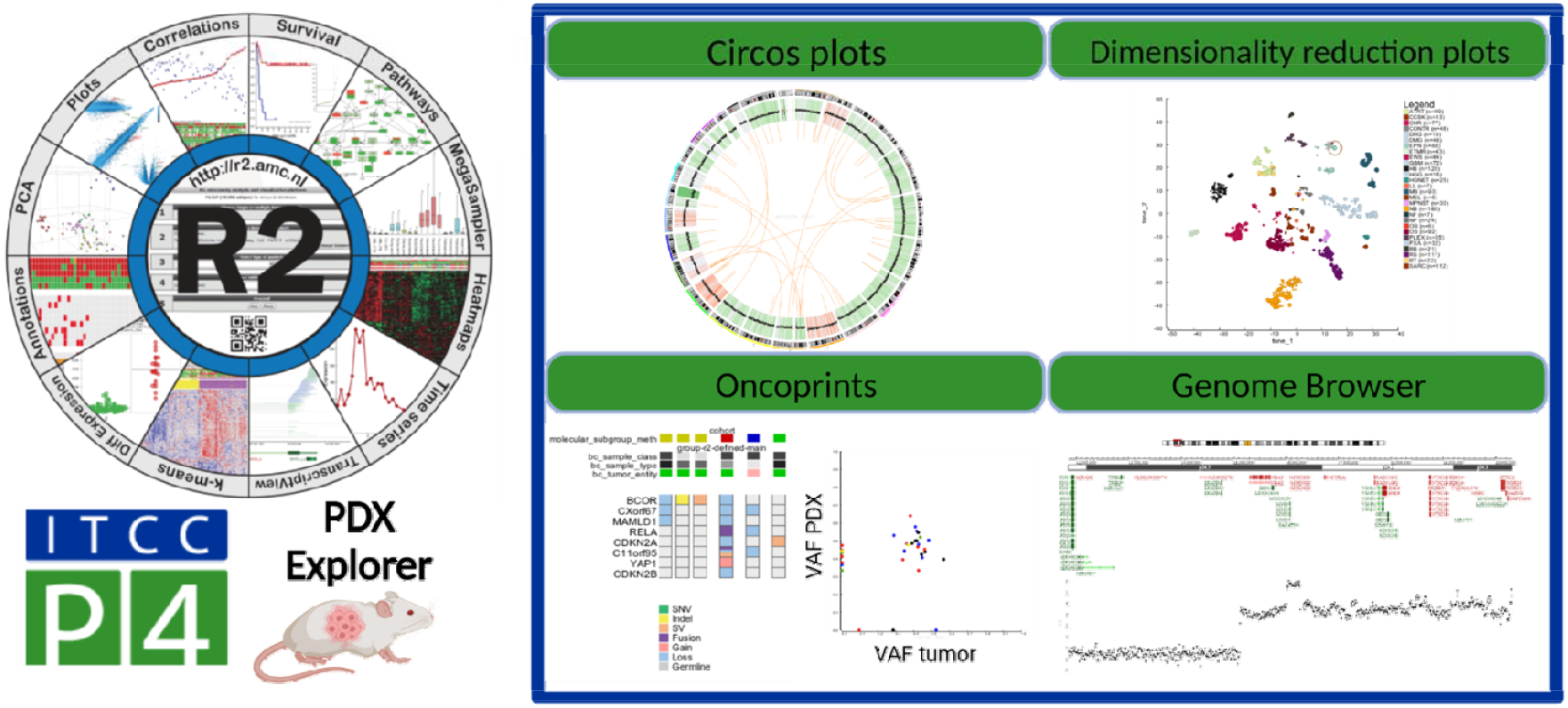
R2 platform-ITCC-P4 PDX datascope. A full report of the clinical information, PDX growth details and molecular characterization for each ITCC-P4 PDX model is available on the ITCC-P4 Datascope in R2 (r2-itccp4. amc.nl). Here, several features were implemented to integrate multi-omic data and to help users in the investigation and visualization of the molecular readouts, such as interactive plots (circos, tSNE, heatmap, etc.), curated oncoprints and the R2 embedded genome browser.

Through the user-friendly customer interface of the portal, registered users can interact with the platform, customize queries, and receive pertinent information tailored to their experimental needs. This user-centric approach enhances data accessibility and promotes a collaborative environment that enables advancing research and therapeutics in the field of oncology.

## Discussion

Significant progress has been made in recent years in comprehending the molecular basis of pediatric tumors. However, to develop impactful therapeutic strategies, there is an urgent need to develop valid and predictive preclinical models for all high-risk childhood cancers and to understand the mechanisms driving treatment resistance.

Among the diverse range of preclinical models, PDX models stand out as particularly suitable for cancer research due to several crucial features. They exhibit overall stability with minimal passages, enabling comprehensive molecular profiling that closely mirrors the genomic, epigenetic, and transcriptomic landscape of the original human tumors. Additionally, under specific conditions, PDX models prove especially valuable for drug testing research, such as orthotopic brain PDXs that offer a surrogate platform for assessing the brain penetrance of deliverable substances. For all these reasons, PDX models have already strongly contributed to preclinical testing (37,38). Yet the use of PDX models in preclinical research also faces constraints, both technical (such as engraftment success rates, immunodeficiency of host mice, and the time-consuming, costly process for model establishment) and biological (including limited availability of certain tumor types and genomic evolution).

The European project “IMI ITCC-P4” focused on establishing an extended and robust PDX resource for pediatric cancer research: the collection of a large and heterogeneous group of established PDX models, with multi-omics molecular data, represents one of the few international and interdisciplinary initiatives created worldwide for childhood cancer modeling.

Recently, several studies have collected childhood tumor PDX models (7,10,18); the PDX resource generated with the MAPPYACTS precision medicine trial (10) included PDX models shared with our ITCC-P4 initiative.

Our study represents a large series of pediatric cancer PDXs with comprehensive genomic characterizations, including the calling of driver mutations, gene fusions and expression profiling, CNVs and DNA methylation profiling. These data demonstrate that the ITCC-P4 PDX repertoire is an excellent representation of the distinct tumor types, subtypes, and molecular vulnerabilities described in pediatric cancer (31). We observed an enrichment of PDX models exhibiting tumorigenic alterations mirroring aggressive phenotypes, like chr1q gain, *CDKN2A/B* loss, *MYC* amplification, *STAG2, TP53* and *ATRX* mutations (39). On the other hand, rare and less aggressive tumor families are still underrepresented, indicating that even more models will be needed. However, less aggressive tumors, such as low-grade gliomas, continue to present considerable challenges for successful engraftment. The ITCC-P4 cohort will be further expanded to include leukemia and lymphoma models, aiming for complete representation of all known molecular subtypes of high-risk pediatric cancers. Evolving the ITCC-P4 cohort, we expand to include leukemia and lymphoma PDX models, while aiming for complete representativity of all known molecular subtypes of pediatric cancers.

A key aspect of our study is that, for the first time, we present molecular characterization of a large set of pediatric tumors PDX models along with the generation, interpretation and comparative analysis of matching patient tumor data and germline controls. This aspect is essential to assess the ability of the xenograft models to recreate the molecular features of the same diseases from which they were generated.

In comparison to previous publications, our PDX molecular dataset now includes DNA-methylome data, which represents a crucial readout for the classification and molecular annotation of several cancer types (31). Methylation-based unsupervised clustering of PDX models resulted in molecular subtype-specific annotations. Also, PDX models segregated according to their entities based on their expression profile and RNA-sequencing data allowed a precise annotation of the frequent, yet essential, fusion events observed in sarcoma and other pediatric tumors. This multi-layered approach was particularly effective in defining the key oncogenic hits for all PDX models, including cases where single omics analyses were insufficient for a comprehensive molecular characterization. In addition to traditional whole-exome data to identify key driver mutations, we generated low-coverage whole-genome sequencing data which enhanced our ability to detect copy number variations and large structural variants. This multi-omics layer integration ensures the accuracy, consistency, and ongoing relevance of specific genetic information (genotyping match and CNV evolution) to obtain a more detailed representation of the molecular status of each PDX model. As the platform expands with the integration of novel PDXs, as well as GEMMs, and organoid models, coupled with advancements in NGS tools and analyses, incorporating additional techniques like ChIPSeq, ATACseq, and DNA and RNA single-cell sequencing become imperative for obtaining a thorough overview of PDX models. These methods play a crucial role in exploring gene regulation and understanding epigenetic modifications and/or cellular diversity between tumor-PDX pairs. Beyond methylation profiling, exploration of the PDX models, and whenever possible, the matched tumor, with regard to histone modifications, chromatin accessibility and nucleosome occupancy, as well as genome organization and long-range genome interactions, will be explored in bulk and at a single-cell level. This will enable the identification of rare cell types and elucidate dynamic changes in gene expression during development, disease progression, or in response to treatments in PDX models.

The molecular comparison between human tumors and matched PDXs frequently demonstrated high concordance regarding genetic variants, tumor mutational burden, and epigenomic and transcriptomic profiles. However, we also observed cases of divergence in the mutational landscape between patient tumor samples and xenografts. Indeed, following correction for tumor cell fractions (TCF) and a normalization method involving the correction of variant allele frequencies (VAF) based on estimated tumor cell purity, an increase or decrease of VAFs upon establishment of PDX could be observed. This suggests either the selection of the observed subclone or the selection of another one. To potentially explain these discrepancies in the enrichment of specific mutations, we considered several factors. First, patient and PDX tumors present different tumor cellularity fractions, with a higher tumor purity in xenografts, while human tumors present the infiltration of additional cellular components coming from the immune system and the microenvironment, which are lost during the establishment of PDX (40). As a result, disparities in tumor cell fractions between matched tumor-PDX pairs can “mask” oncogenic hits in highly infiltrated tumor samples, while these drivers may become detectable in the corresponding tumor-cell enriched PDX data. Additionally, somatic mutations predominantly enriched in either PDX or tumor samples may result from subclonal selection during xenograft establishment or the absence of the immune system, providing a growth advantage for specific variants in the PDX model (35).

A further strength of our collection of PDX models is the inclusion of a subset of “serial” models generated from patient tumors collected at different phases over the course of disease. The characterization of these models highlighted the importance of investigating the molecular changes that occur across multiple tumor recurrences and/or upon exposure to treatments. This analysis might represent a crucial step to gaining a better understanding of pediatric tumor progression, evolution, and treatment resistance. Our findings may also help to identify alternative treatment options for resistant recurrent tumors or metastases.

Even though molecular analyses show that most PDX models show close overlap with the matched human tumors with regards to molecular features, some cellular differences may so far have remained undetected. Bias may also arise from the site of engraftment, as in this cohort of PDX models, non-brain tumors were engrafted subcutaneously and not in the same anatomical location as the primary tumor. However, the overall resource is of crucial importance, as the data generated within this study will serve as a strong basis for further investigation in the field of pediatric cancer research, particularly for the identification and validation of novel therapy options based on molecular vulnerabilities through preclinical *in vivo* drug testing on these PDXs. The drug sensitivity results collected so far from the ITCC-P4 PDX models aligned with their molecular status, supporting the importance of performing proof-of-concept drug testing studies.

This approach, extended to the full repertoire of models generated within the ITCC-P4 platform, represents the core service provided by the non-for-profit company ITCC-P4 gGmbH (gGmbH designating a company with limited liability according to German law) in close collaboration with three contract-research organizations (CROs), with the aim of supporting drug testing and research for pediatric cancer, ITCC-P4 GmbH Paediatric Preclinical Proof of Concept Platform). In this unique organization, ITCC-P4 and the CROs do not distribute or sell any of the PDX models, but importantly, the PDX models are accessible for preclinical drug testing for academic or industrial users via the non-profit ITCC-P4 gGmbH company, with the CROs performing the actual drug testing or experimental approaches, while the models are co-owned by the academic partners. Therefore, all PDX models are accessible and can be shared for scientific collaborations via the academic partners in the consortium. In addition to methylation profiling, PDX models via the ITCC-P4 gGmbH provide advantages, compared to purely academic collaborations, as the ITCC-P4 gGmbH mouse trials include tumor engraftment, treatment, tumor response monitoring, and data analyses and reporting. Furthermore, the possibility of choosing between different models depending on tumor type and subtype, and molecular features, and the fully established workflow within the ITCC-P4 gGmbH are of interest to the scientific community given the possibility of a rapid set-up and completion of these experiments. In addition, engaging the specialized CROs embedded in ITCC-P4 gGmbH for *in vivo* studies ensures the highest quality standards and ethical compliance, fostering adherence to the 3Rs principle.

The ITCC-P4 PDX Data scope in R2, collecting the molecular data generated for the ITCC-P4 PDX models (and matched patient tumors), will be accessible to the whole scientific community, making this platform a unique resource for the field.

In conclusion, the ITCC-P4 platform represents a valid and powerful tool to investigate the biology of pediatric cancer based on the establishment and molecular characterization of pediatric cancer PDX models, contributing in this way to the development of innovative therapeutic options for childhood cancer patients.

## Methods

### Patient sample collection

Patient-derived material and clinical data were collected under institutional ethical approval (University of Heidelberg and collaborating centers) and in accordance with the Declaration of Helsinki. Informed consent was obtained from all participants or their legal guardians. Tumor and matched germline samples were collected as part of the MAPPYACTS study and collaborating ITCC-P4 sites. Detailed information on patient enrollment, sample collection, consent procedures, and data management is provided in the Supplementary Methods

### PDX model establishment

All animal procedures complied with institutional and European guidelines (Directive 2010/63/EU) and were approved by local ethical committees and national authorities (France and Italy). Fresh tumor tissues from MAPPYACTS patients were implanted into immunodeficient mice to generate patient-derived xenograft (PDX) models. Orthotopic models were established for brain tumors and subcutaneous models for non-brain tumors, following approved analgesia and anesthesia protocols. PDX tumors were expanded across passages and cryopreserved for molecular characterization and biobanking. Detailed procedures for tumor dissociation, transplantation, animal care, monitoring, and reimplantation are described in the Supplementary Methods.

### Nucleic acid extraction

DNA and RNA were extracted from tissue samples using automated purification systems and standardized protocols. Nucleic acid yield and integrity were assessed by fluorometric quantification and electrophoretic analysis, with RNA samples meeting a minimum integrity threshold (RIN ≥ 7). Details of extraction kits, instrumentation, and quality control procedures are provided in the Supplementary Methods.

### ITCC-P4 barcoding system

Human and PDX samples, along with associated clinical and molecular data, were anonymized and tracked using an automated barcoding system. Details of the coding structure and data registration process are provided in the Supplementary Methods.

### Molecular profiling and data processing

#### Whole genome and exome sequencing workflow

Patient and PDX tumor samples underwent library preparation and sequencing at two ITCC-P4 partner sites (DKFZ and Institut Curie) using low-coverage whole-genome (lcWGS) and whole-exome sequencing (WES). Additional raw sequencing data were obtained from collaborating institutions. Sequencing reads were aligned to a combined human–mouse reference genome, and variant calling was performed using standardized in-house pipelines. Full details of library preparation, sequencing platforms, and bioinformatic workflows are provided in the Supplementary Methods.

#### Processing Samples without Patient-Matched Germline Data: The “No-control workflow”

For PDX samples lacking matched germline controls, variant detection was performed using an established in-house “No-Control” workflow integrating multiple variant-calling tools. Common germline variants were excluded using population and reference databases, and functional annotation and filtering were conducted to prioritize rare, potentially pathogenic variants. Detailed pipeline configurations and filtering parameters are provided in the Supplementary Methods.

#### DNA methylation profiling and analysis

Genomic DNA from patient tumors and matched PDX samples was profiled using Illumina Methylation450K and EPIC BeadChip arrays at DKFZ and Institut Curie. Data processing, normalization, and quality control were performed using the RnBeads Bioconductor package. Tumor classification and molecular subtype assignment were based on established DNA methylation classifiers and unsupervised clustering analyses. Full preprocessing parameters, filtering criteria, and classifier references are described in the Supplementary Methods.

#### Copy number variation analysis

CNVs in patient tumors and PDX models were characterized using multiple complementary computational tools, including ichorCNA, Sequenza, CNVkit, and conumee, to ensure accurate detection of large-scale and focal copy number alterations and reliable tumor purity estimation. Sequencing- and methylation-based approaches were combined to provide a comprehensive CNV profile. Full details of pipelines, parameters, and input requirements are provided in the Supplementary Methods.

#### RNA sequencing and analysis

RNA sequencing of patient tumors and PDX models was performed on Illumina HiSeq instruments. Data were processed using the established DKFZ OTP in-house RNA-seq workflow (based on the Roddy framework), including read alignment with STAR, gene expression quantification with FeatureCounts, and gene fusion detection with Arriba. Processed expression data were used for unsupervised clustering, differential gene expression, and gene ontology analyses. Full details of pipeline versions, parameters, and downstream analyses are provided in the Supplementary Methods.

### Identification and Annotation of Driver Genes for Mutational Landscape Analysis

Mutational landscape and variant analysis Driver genes were identified and annotated using curated references, including the 2022 WHO Classification of Pediatric Tumors and published reviews on pediatric solid cancers. Tumor mutational burden (TMB) was calculated from high-confidence coding SNVs and indels derived from whole-exome sequencing. Variant allele frequencies (VAFs) were computed to assess intra-tumor heterogeneity and clonal evolution in PDX models. PDX–tumor correlations were further evaluated using metrics such as delta-VAF and VAF ratios. Statistical analyses, including correlation and fold-change calculations, were performed in R. Full workflows, formulas, and analytical parameters are provided in the Supplementary Methods.

## Figure legend

**Supplementary Figure 1.**
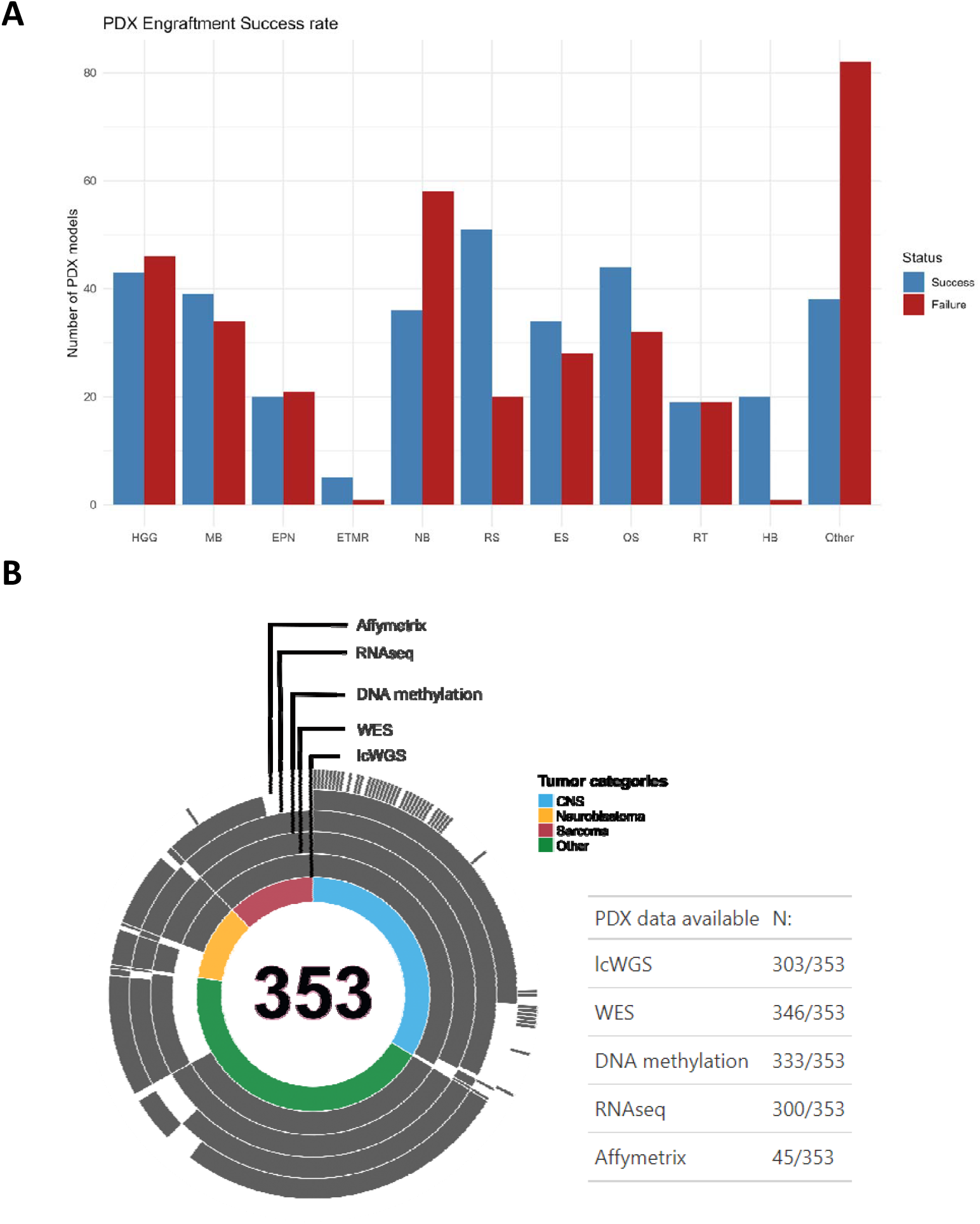
A) Barplot showing the number of successfully vs. failed engrafted PDX models within the ITCC-P4 platform and representing the main pediatric cancer entities. B) Resume of multi-omic data coverage for the whole ITCC-P4 PDX cohort described in this study.

**Supplementary Figure 2.**
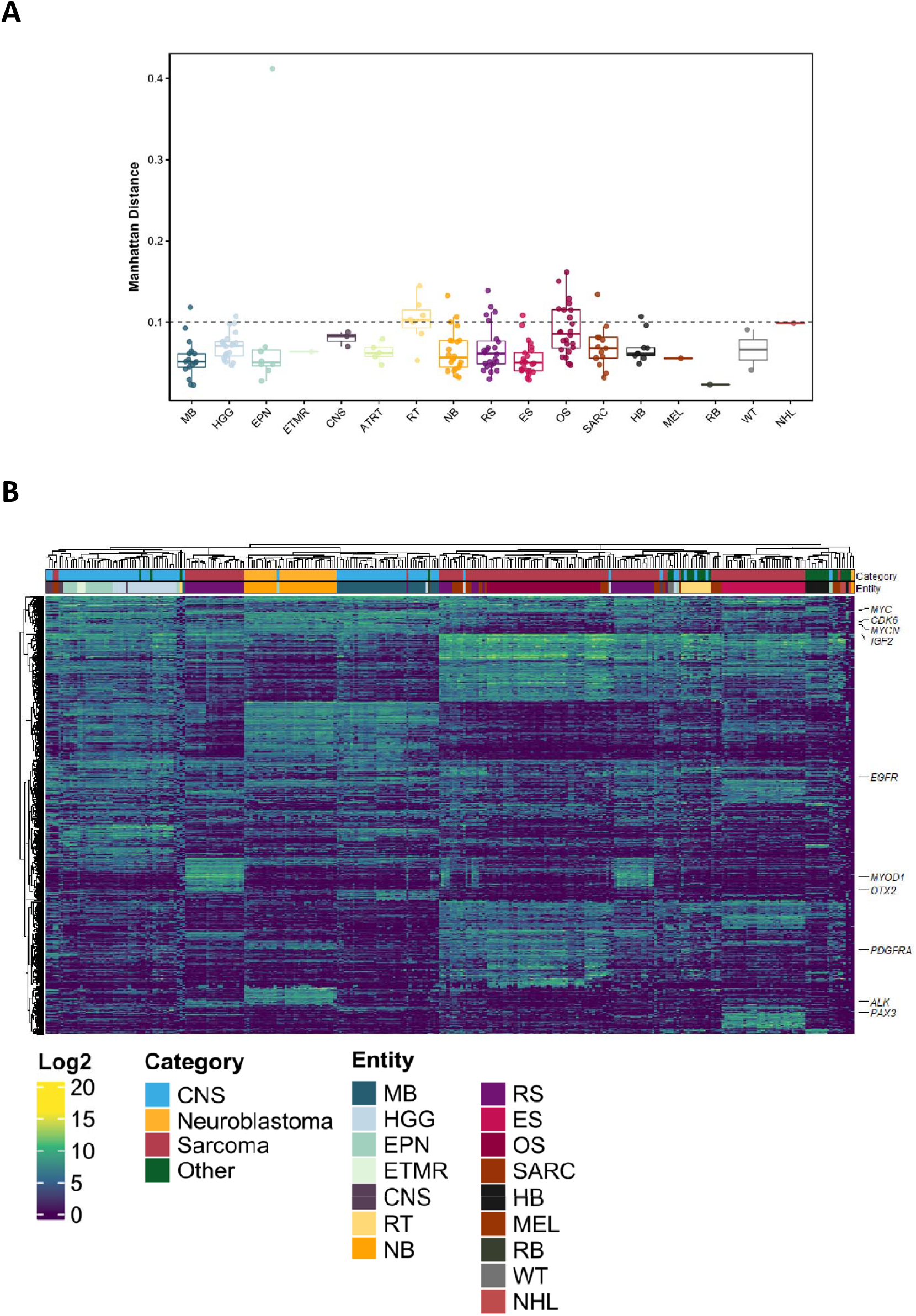
A) PDX-tumor Manhattan distance measures of the methylation signals overlapping SNPs. Samples were grouped according to the tumor entities (assigned based on the methylation tumor classifier methods) represented in the ITCC-P4 dataset B) Heatmap showing the unsupervised hierarchical clustering analyses of ITCC-P4 PDX models based on the gene expression values (expressed in Log2 scale) of the top 1000 most variable (SD) expressed genes across the cohort. The expression of several key gene markers for a variety of pediatric tumor entities was highlighted in the plot.

**Supplementary Figure 3.**
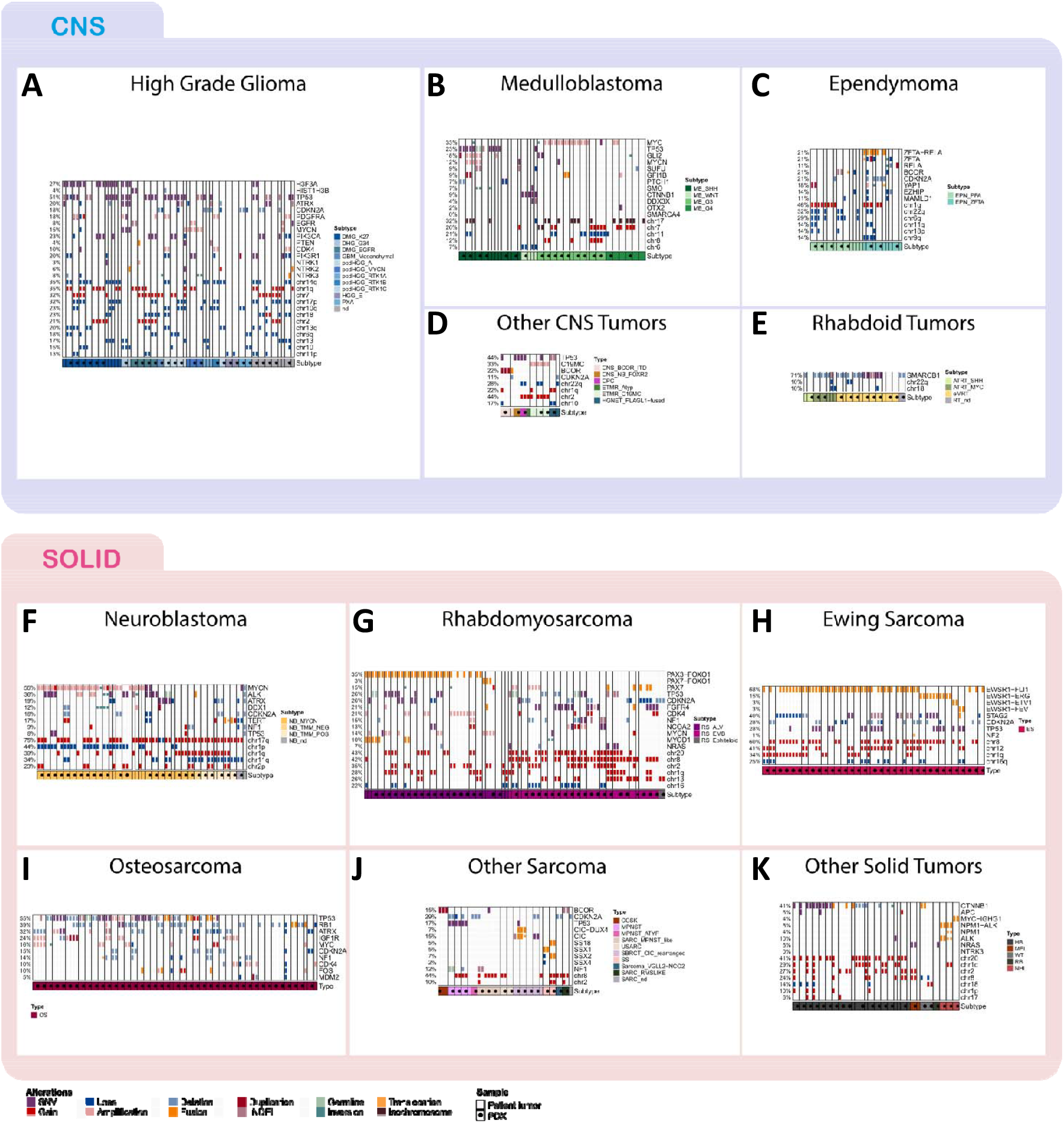
Comparative overview of the mutational landscape of ITCC-P4 PDX models (black dots) and matched tumor samples. Each panel resumes the driver mutations observed in PDX and patient data for the main tumor types present in this study: High-grade glioma (A); Medulloblastoma (B); Ependymoma (C); Other CNS tumors (D); Rhabdoid tumors (E); Neuroblastoma (F); Rhabdomyosarcoma (G); Ewing sarcoma (H); Osteosarcoma (I); other sarcomas (J); other solid tumors (K). Tumor subtypes were indicated for each sample, if known and defined based on the methylation analyses.

**Supplementary Figure 4.**
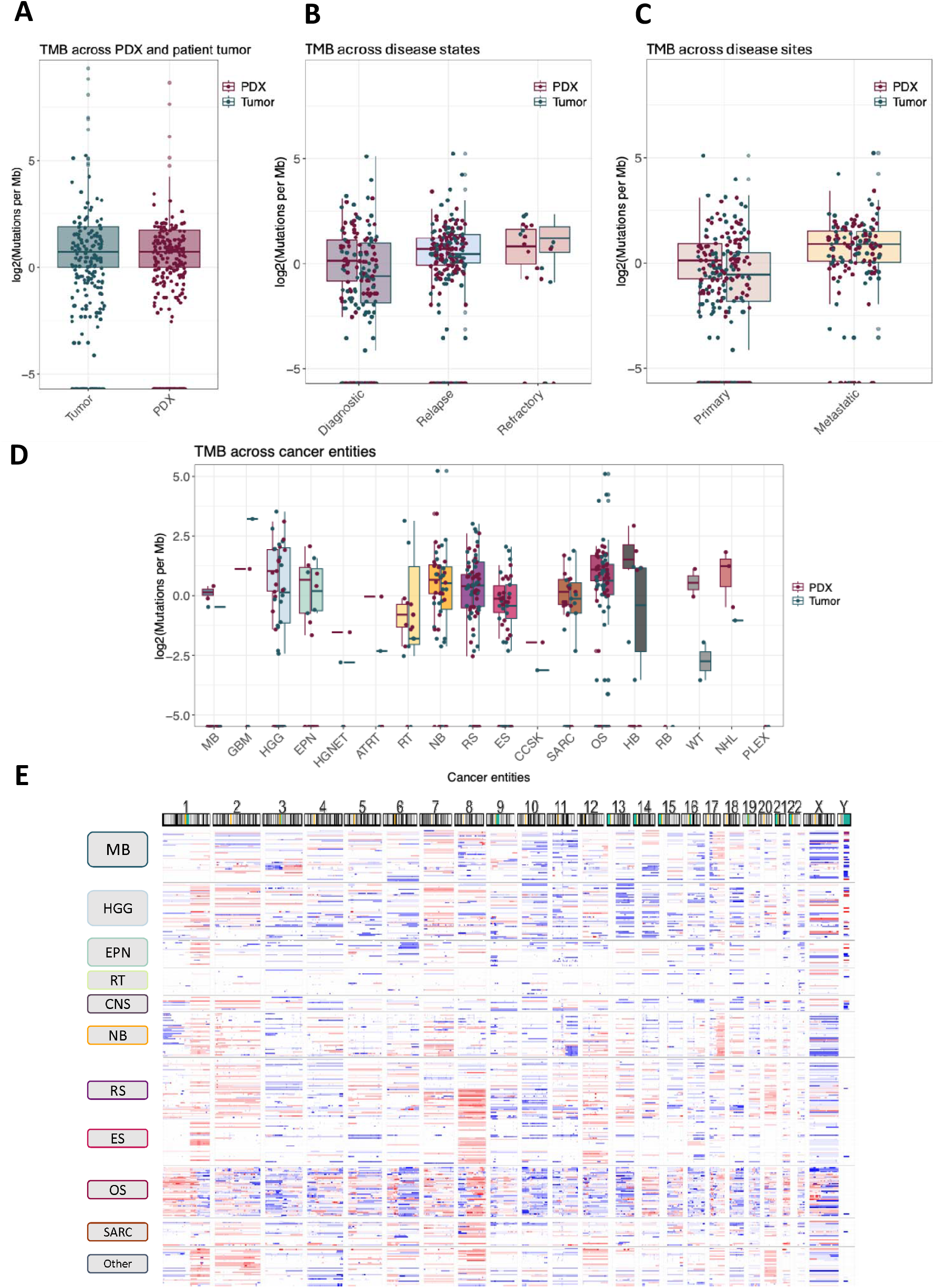
A) Tumor mutational burden (TMB; log2 scaled values) of PDX models (red) and patient tumors (blue); B) TMB values of PDXs (red dots) and human tumors (blue dots), grouped according to the disease state; C) TMB values of PDXs (red dots) and human tumors (blue dots), grouped according to the disease site; D) TMB values of PDXs (red dots) and human tumors (blue dots), grouped according to the tumor types; E) Heatmap showing the whole chromosome CNV landscape of the ITCC-P4 PDX models, grouped according to their tumor types. Gains of chromosome regions are shown in red, while region losses are represented by blue segments.

**Supplementary Figure 5.**
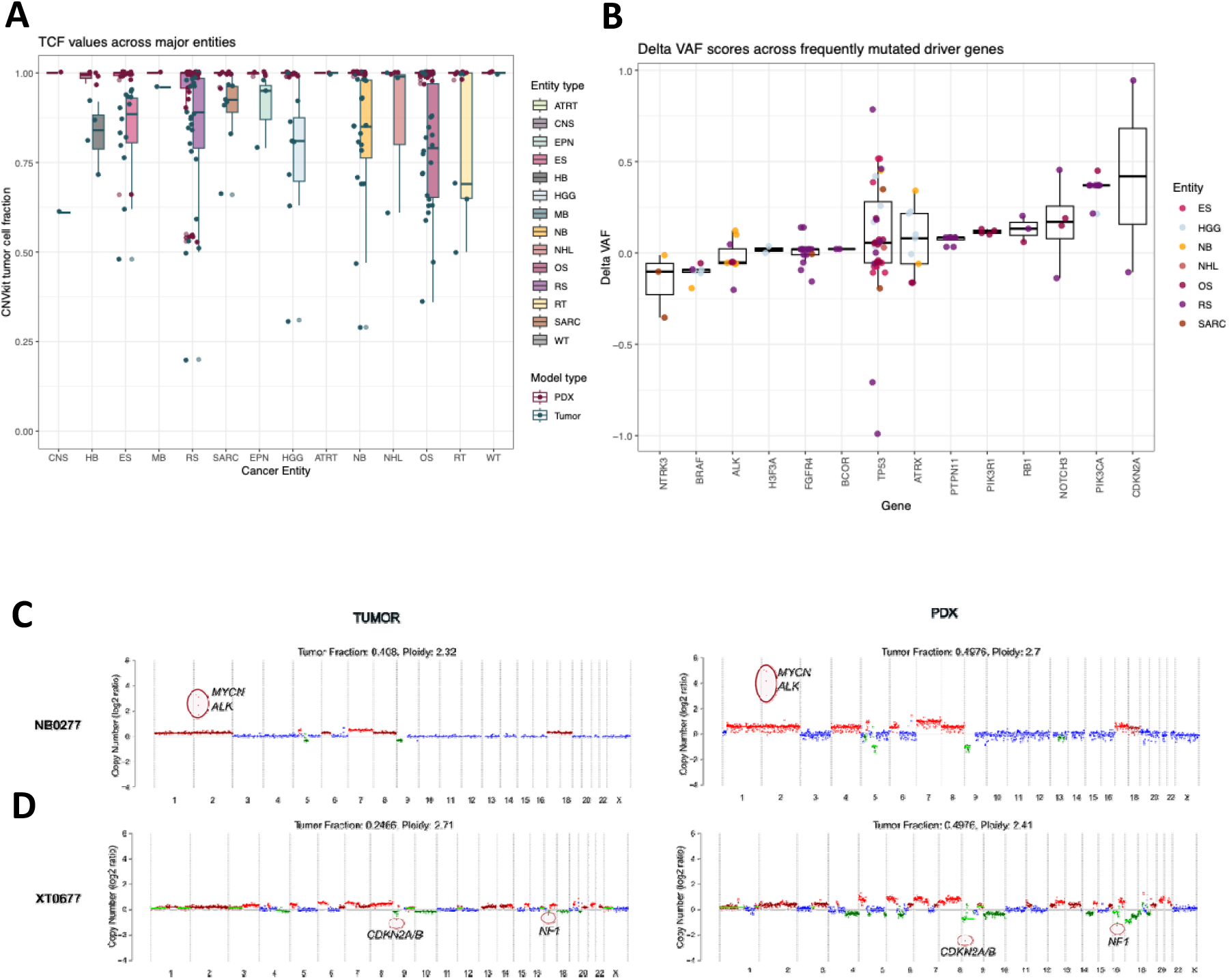
A) Tumor cellular fraction (TCF), calculated using CNVkit for each PDX (red dots) and patient (blue dots) sample within the ITCC-P4 dataset and grouped according to their tumor type; B) Delta VAF scores of the most frequently mutated genes in the ITCC-P4 dataset. Dot colors refer to the tumor type defined for each PDX/tumor pair; C) CNV plots of model NB0277 (right) and relative patient tumor (right). The focal amplification of MYCN and ALK genes (red circles) could be detected in both cases; D) CNV plots of XT0677 PDX (right) and original tumor (left). The chromosome and focal gene alterations, including *CDKN2A/B* deletion and NF1 loss, observed in the two plots are overlapping;

**Supplementary Figure 6.**
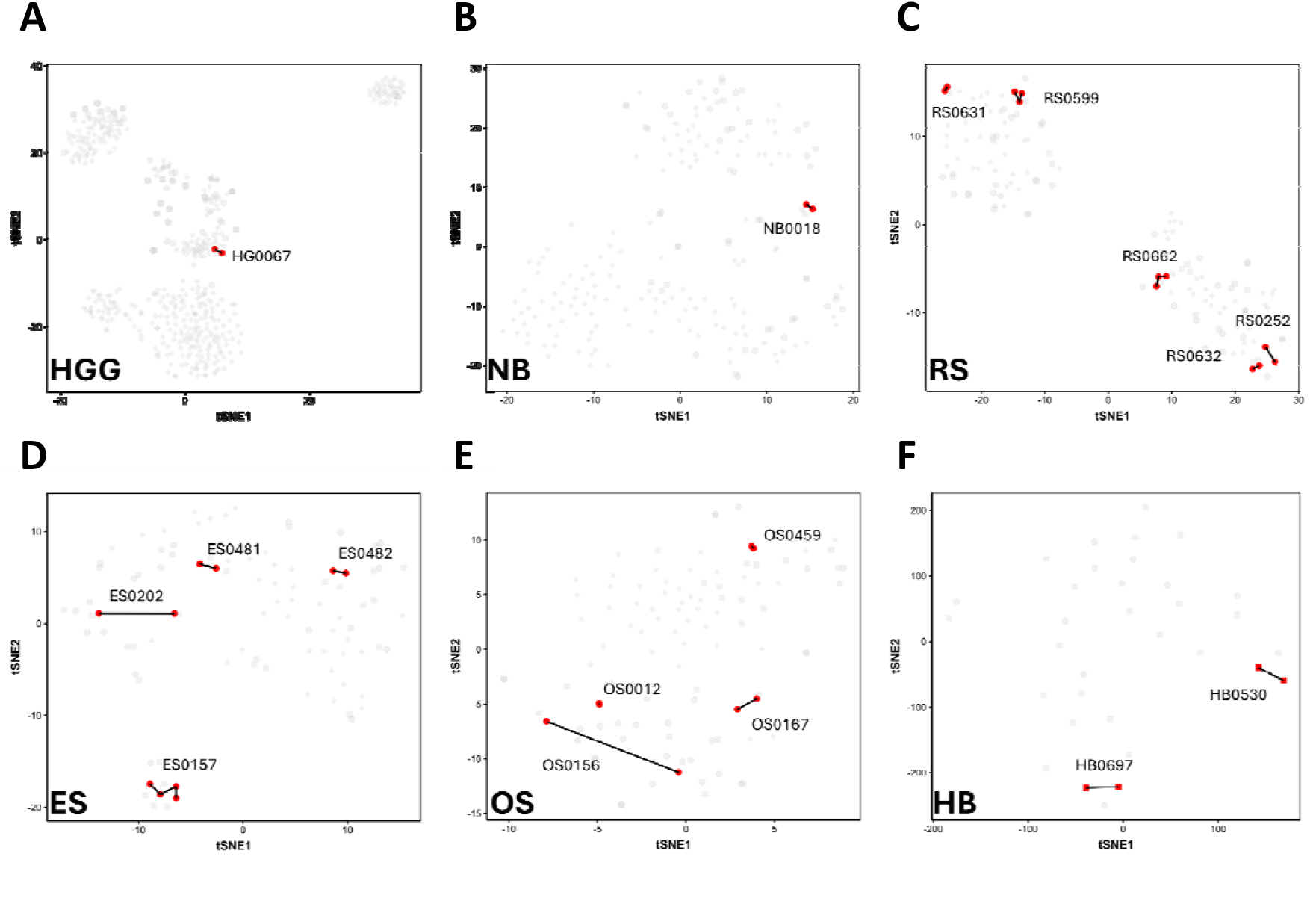
A-F) tSNE plots of high-grade glioma (A), neuroblastoma (B), rhabdomyosarcoma (C), Ewing sarcoma (D),osteosarcoma (E) and hepatoblastoma (F) methylation samples (as already shown in figures 2 and S2), where serial PDX models are highlighted (red dots) and connected (black lines). F) CNV plots of RS0252 (primary and relapse) PDX models. Key focal events are highlighted in red circles.

## Data availability statement

The curated multi-omics data are accessible via the R2 ITCC-P4 PDX Datascope portal (http://r2platform.com/itcc-p4/). Raw sequencing data are accessible via EGA (access number: EGAS00001008330).

Any other data shown in this study are available upon request. For inquiries regarding the data, please contact ITCC-P4 gGmbH.

## Acknowledgments

*ITCC-P4 IMI-2 project:*

Funding sources/grant numbers: “This project has received funding from the Innovative Medicines Initiative 2 Joint Undertaking under grant agreement number 116064. This Joint Undertaking receives support from the European Union’s Horizon 2020 research and innovation program and EFPIA”. The project was supported by funding from Bayer, Roche, Servier within the IMI2 ITCC P4 program. Molecular characterization was supported by funding from Roche, Pfizer, Amgen within the IMI2 ITCC P4 program.

*INFORM/DKFZ, Germany:*

The INFORM program is financially supported by the German Cancer Research Center (DKFZ), several German health insurance companies, the German Cancer Consortium (DKTK), the German Federal Ministry of Education and Research (BMBF), the German Federal Ministry of Health (BMG), the Ministry of Science, Research and the Arts of the State of Baden-Württemberg (MWK BW); the German Cancer Aid (DKH), the German Childhood Cancer Foundation (DKS), RTL television, the aid organization BILD hilft e.V. (Ein Herz für Kinder) and the generous private donation of the Scheu family.

We would like to express our sincere thanks to Carsten Maus, Erjia Wang (Genomics and Proteomics Core Facility, DKFZ). Gregor Warsow (Omics IT and Data Management Core Facility, DKFZ) for their highly dedicated support in data management and processing. Gnanaprakash Balasubramanian and Rolf Kabbe (Division of Pediatric Neurooncology, DKFZ) for their sincere and dedicated contribution to the bioinformatics analyses.

*Institut Curie, France:*

Resources/Facilities: At Institut Curie, this work was supported by Imagine for Margo, the Annenberg Foundation, the Association Hubert Gouin Enfance et Cancer, the Fédération Enfants Cancers Santé, the Société Française de lutte contre les Cancers et les leucémies de l’Enfant et l’adolescent (SFCE), Les Bagouz à Manon, Les amis de Claire, La Ligue contre le Cancer, and the Fondation ARC pour la Recherche contre le Cancer (ARC). Funding was also obtained from INCa/SiRIC/ (Grant INCa-DGOS-4654), and PRTK2019-1-RT-02-ICR-1 grant. High-throughput sequencing was performed by the ICGex NGS platform of the Institut Curie supported by the grants ANR-10-EQPX-03 (Equipex) and ANR-10-INBS-09-08 (France Génomique Consortium) from the Agence Nationale de la Recherche (“Investissements d’Avenir” program), by the Canceropole Ile-de-France and by the SiRIC-Curie program - SiRIC Grant “INCa-DGOS-4654”. Tumor sequencing data was also contributed from the MAPPYACTS protocol (clinicaltrial.gov: NCT02613962) and MICCHADO (NCT03496402). The authors thank Angela Bellini and Rachida Bouarich for their contributions.

*ACC, Italy:*

Funding sources/grant numbers: Associazione Bianca Garavaglia (ABG) A/15/01N and A/18/01A

IRCCS—Istituto Ortopedico Rizzoli, Experimental Oncology Laboratory, Bologna, Italy, (Katia Scotlandi, Maria Cristina Manara, Lorena Landuzzi) the study was supported by the Innovative Medicines Initiative 2 Joint Undertaking ITCC P4 under grant agreement number 116064.

*Institute Gustave Roussy, France:*

At Gustave Roussy the work was supported by grants from Fondation Gustave Roussy; Fédération Enfants Cancers et Santé, Société Française de lutte contre les Cancers et les leucémies de l’Enfant et l’adolescent (SFCE), Association AREMIG and Thibault BRIET; Parrainage médecin-chercheur of Gustave Roussy; INSERM; Canceropôle Ile-de-France; Ligue Nationale Contre le Cancer (Equipe labellisée); Fondation ARC for the European projects ERA-NET on Translational Cancer Research (TRANSCAN 2) Joint Transnational Call 2014 (JTC 2014) ‘Targeting Of Resistance in PEDiatric Oncology (TORPEDO)’.

*Xentech, France:*

PDXs were established with the Paris hospitals’ network in France in collaboration with Sophie Branchereau, Pediatric Surgery Department, Bicêtre Hospital, Le Kremlin Bicêtre, France. RNAseq and WGS fastq files, of Xentech models, were generated by Carolina Armengol, Childhood Liver Oncology Group (c-LOG), Health Sciences Research Institute Germans Trias i Pujol (IGTP), Badalona, Liqin Zhu, Department of Pharmacy and Pharmaceutical Sciences, St. Jude Children’s Research Hospital, Memphis, TN, USA.

*UZH, Switzerland:*

At UZH, this work was supported by Swiss Cancer League projects KLS-5143-08-2020 and KFS-5422-08-2021, the Clinical Research Priority Program (CCRP) “Precision Heamatology/Oncology” and the Childhood Cancer Research Foundation Switzerland, the Swiss National Science Foundation projects 3100-175558 and 10.000.473, the Balgrist Foundation, the ResOrtho Foundation, the FORCE Foundation, the Bryn Turner-Samuels Foundation, the Pierre Mercier Foundation and the Sarcoma Foundation of America. We would like to express our sincere thanks to Jean-Pierre Bourquin, Beat Bornhauser, Irina Banzola, Stephanie Kasper, Willemijn Breunis, Daniel Müller, Sander Botter, Knut Husmann and the Swiss Center for Musculoskeletal Biobanking (SCMB).

*EPO, Germany:*

Funding sources/grant numbers: “This project has received funding from the Innovative Medicines Initiative 2 Joint Undertaking ITCC P4 under grant agreement number 116064.

Individuals/Organizations contributing to this work: Svetlana Gromova (technical assistance)

Resources/Facilities: The project has been performed at the facilities and using Resources of EPO Experimental Pharmacology & Oncology Berlin-Buch GmbH.

*CHARITÉ, Germany:*

Funding sources: This project has received funding from the Innovative Medicines Initiative 2 Joint Undertaking ITCC P4 under grant agreement number 116064. This work was further supported by the German Cancer Consortium (DKTK) and the parent’s initiative at Charité KINDerLEBEN e.V.

*ICR, United Kingdom:*

At ICR, this work was supported by grants to Chris Jones from Cancer Research UK (DRCRPG-Nov21\100002), Abbie’s Army, Cris Cancer Foundation, the Ollie Young Foundation and Lucas’ Legacy. We acknowledge NHS funding to the ICR/Royal Marsden Hospital Biomedical Research Centre. At ICR, Prof Louis Chesler is financially supported by UK HEFCE/ICR support. We thank the patients, their families and the staff at the Royal Marsden NHS Foundation Trust, Great Ormond Street Hospital for Sick Children and the Royal Manchester Children’s Hospital for their support in participation and donation of critical samples for this research. Prof Chesler and Dr Tucker would personally like to acknowledge Dr Paola Angelini and Mr Tony Rogers for their vital help in the delivery of this project at ICR.

*FSJD, Spain:*

At Hospital Sant Joan de Deu we acknowledge the clinical fellows involved in the care of patients and the families of the children for donating the samples for research. We thank the Xarxa de Bancs de Tumors de Catalunya (XBTC; sponsored by Pla Director d’Oncologia de Catalunya).

*MUW, Austria:*

University of Vienna: City of Vienna Fund for Innovative Interdisciplinary Cancer Research (#21165)

*Charles University and University Hospital Motol, Czech Republic:*

Ministry of Health of the Czech Republic, MH CZ–DRO, University Hospital Motol, Prague, Czech Republic (00064203)

*Others:*

We thank the Pediatric brain tumor research fund guild Run of Hope for their support in creating the PDX models described in this study.

We thank Olaf Heidenreich, Elizabeth Schweighart, Jean-Pierre Bourquin, Beat Bornhauser, Irina Banzola for their contribution to the ITCC-P4 consortium.

## Competing interests

The authors declare that they have no competing interests related to this study/ Alternatively, specify any potential conflicts of interest.

Stefan Pfister, Co-founder and shareholder Heidelberg Epignostix GmbH

Natalie Jäger is a full-time employee of Heidelberg Epignostix GmbH

Martin Sill, Co-founder and shareholder Heidelberg Epignostix GmbH

Jens Hoffmann: Shareholder EPO Experimental Pharmacology & Oncology Berlin-Buch GmbH

Justyna Wierzbinska and Andreas Schlicker are employees of Bayer AG. Andreas Schlicker is a shareholder of Bayer AG.

Petra Hamerlik provides consultancy for LindonLight Collective and Rakobina Therapeutics.

Stefano Cairo is now a full-time employee of Champions Oncology, Rockville, Maryland, USA

David Shields is an employee of Pfizer Inc and holds shares in the company.

Maureen M. Hattersley is an employee of AstraZeneca and holds shares in the company.

Employees from the following pharmaceutical companies also contributed as co-authors to the ITCC-P4 consortium project, as stated in their affiliations: LILLY, ROCHE, PFIZER, BAYER,PHARMA MAR, CHARLES RIVER, JANSSEN, AZ, AMGEN, SERVIER, SANOFI.

