## Supplemental Methods for "The ITCC-P4 PDX platform of pediatric cancers for preclinical testing"

**Supplementary Methods**

**Patient sample collection**

The study was approved by the Ethics Board of the Medical Faculty of the University of Heidelberg or local institutions. We obtained informed consent from all patients or their parents/caregivers for the collection of biological samples, including permission for anonymized data sharing in academic and non-profit settings, performed according to the Helsinki guidelines. All models developed within the MAPPYACTS study had been developed following informed consent for this study from patients and legal representatives [(1,2)](https://sciwheel.com/work/citation?ids=15356496,15379858&pre=&pre=&suf=&suf=&sa=0,0). For each patient, fresh tumor material was collected (by surgical resection, or biopsy), as well as a constitutional (germline) sample (mainly the patients´ blood sample), when available. Additional models established by non-ITCC-P4 sites were included via direct collaboration with the ITCC-P4 consortium. Clinical information (age at diagnosis; gender; tumor location; tumor stage, tumor characteristics, such as primary or metastatic site, diagnostic or relapse sample; pathological and molecular diagnosis report; treatment history; basic follow-up information) were collected if available.

**PDX model establishment**

Animal care and use were performed in accordance with international guidelines and the recommendations of the European Community (2010/63/UE). Experimental procedures within MAPPYACTS were specifically approved by the ethics committee, the France Ministry of Agriculture or Italian Ministry of Health; Gustave Roussy CEEA26 (CEEA PdL N°6, approval number: 2015032614359689 V7, 1281.01, C75-05-18, 2012-017), Institut Curie CEEA-IC #118 (APAFIS#11206-2017090816044613-v2), Fondazione IRCCS Istituto Nazionale dei Tumori (OPBA authorization: INT 03_2018, Italian Ministry of Health authorization: 646/2018-PR). Fresh tumor samples transferred in cell culture medium (Neurobasal, NeuroCult NS or RPMI 1640, without FCS or other growth factors) were immediately transplanted into NSG (NOD.Cg-Prkdcscid IL2rgtm1Wjl/SzJ), Swiss Nude (Crl:NU(Ico)-Foxn1nu) or CB17 SCID (CB17/Icr-Prkdcscid/IcrIcoCrl) immunodeficient mice. Each PDX model was considered to be established if a tumor has grown in the P2 mice and if a revived serum/DMSO frozen tumor fragment can regenerate the PDX model.

*Orthotopic injections of brain tumors*

Fresh brain tumor tissue was dissociated into a single cell suspension by careful pipetting or by treatment with accutase (Accumax, eBioscience) and incubated for 15 min at 37 °C. Next, the cell suspension was centrifuged at 1200 rpm, the supernatant was carefully removed, and 1 ml cell culture medium was added again before cells were pushed through a cell strainer (Neolab). Next, the cells were counted and assessed for viability; ~ 2x10^6^ cells were stored for molecular characterization, while additional ~1x10^6^ cells were intracranially injected into 2-6 NSG mice. For perioperative analgesia, mice are subcutaneously injected with carprofen (5 mg/kg). Twenty minutes later, animals are anesthetized by isoflurane inhalation (1.5-2.5Vol%). After negative reflex testing, animals were fixated in a stereotactic frame and a 5 mm incision in the scalp was introduced. For local anesthesia, 0.25% bupivacaine was applied to the incision. An 18G cannula (diameter: 1.27 mm) was used to drill a small hole into the skull. Next, the Hamilton needle was introduced into the brain within specific injection sites, depending on the original location of the human tumor to inject the cell suspension. Following the operation, mice were constantly monitored and treated with carprofen (5 mg/kg) for the following 1-7 days in case of pain symptoms. As a follow-up, mice were regularly observed for neurological or other tumor-related symptoms. Tumor growth was monitored by MRI. When the brain tumor reached the maximum volume and for subsequent passages (P1-2), tumors were removed from euthanized mice and again dissociated into a single-cell suspension. Snap-frozen pellets of at least 1x10^6^ cells were collected for molecular characterizations, and cryovials of 1.5x10^6^ cells (in 90% Neurobasal or NeuroCult NS medium and 10% DMSO) were collected for biobanking and model propagation. while the remaining material was split into cryovials (1.5x106 cells per vial, 90% Neurobasal or NeuroCult NS medium / 10% DMSO).

*Subcutaneous injections of non-brain tumors*

Fresh non-brain tumors were dissected into several small fragments (typically for 5 to 100mm^3^, depending on the size of the tumor) and transplanted into the flank or under the intrascapular fat pad of 2 – 6 NSG mice, anesthetized either by isofluorane inhalation or by intraperitoneal injection of a mixture of Xylazine (Rompun® 5 to 10 mg/kg) and Ketamine (Imalgene® 80 to 100 mg/kg). The mice were regularly observed for tumor growth, and tumors were measured at constant intervals during their growth. For subsequent passages, the tumors were removed from euthanized mice and dissected in a sterile cell culture dish into several small fragments (typically 20 to 100 mm^3^, depending on the size of the tumor).

**Nucleic acid extraction**

DNA and RNA were isolated from tissue samples using the automated Maxwell nucleic acid purification system and the respective kits for either blood or tissue samples., or equivalent standardized procedures [(1)](https://sciwheel.com/work/citation?ids=15356496&pre=&suf=&sa=0). DNA quantity and quality were measured using Qubit and high-resolution electrophoresis. RNA quantity and quality were measured using the Agilent Bioanalyzer system. The RNA Integrity Number (RIN) values threshold was set as ≥ 7.

**ITCC-P4 barcoding system**

All human and corresponding PDX samples (cryo vials; cell pellets; DNA; RNA), as well as clinical, PDX (tumor growth latency, tumor penetrance) and molecular data were anonymized and labeled using a barcoding system based on the original source, tumor entity, tumor event, sample type and data type. Barcodes were automatically generated once samples and patient data were registered through the online system in R2 (r2-itccp4. amc.nl).

**Molecular profiling and data processing**

**Whole genome and exome sequencing workflow**

Patient and PDX tumor samples were subjected to library construction followed by sequencing at two ITCC-P4 partner sites (DKFZ and Institut Curie) employing the Paired-End Sequencing method (lcWGS, and WES), as described previously [(1,3)](https://sciwheel.com/work/citation?ids=15356496,11559627&pre=&pre=&suf=&suf=&sa=0,0). Additionally, we acquired raw NGS data (fastq files) from various collaborating institutions, which included low-coverage whole genome sequencing (lcWGS) and whole exome sequencing (WES) data. These samples were independently profiled, resulting in the use of different library preparation kits and sequencing platforms depending on whether it was WES (**Table 1**) or WGS (**Table 2**).

| **Institution** | **Library Preparation kit** | **WES Sequencing platform** |
| --- | --- | --- |
| DKFZ | Agilent SureSelectXT HS | Illumina HiSeq 2000 |
| CCIA | TruSeq Nano DNA HT | Illumina HiSeq X Ten |
| CURIE | Agilent SureSelect Clinical Research Exome V2 | Illumina NovaSeq 6000 |
| IGR | Agilent SureSelect Clinical Research Exome V2 | Illumina NextSeq 500 |
| ICR | Agilent SureSelect Clinical Research Exome V7 | Illumina NovaSeq 6000 S2 |

*Table 1 : WES Library Preparation Kits and Sequencing Platforms Utilized in ITCC-P4 Study*

| **Institution** | **WGS Sequencing platform** |
| --- | --- |
| DKFZ | Illumina NovaSeq 6000 S1 |
| CCIA | Illumina HiSeq X Ten |
| CURIE | Illumina NovaSeq 6000 |
| Xentech | Illumina HiSeq 2500 |

*Table 2:* lcWGS Library Preparation Kits and Sequencing Platforms Utilized in ITCC-P4 Study

The sequencing reads of the patient tumor and PDX samples, with corresponding germline controls if available, were aligned to a merged reference genome, which combined the human (hs37d5) and murine (GRChm38mm10) reference genomes. This alignment was performed using the BWA alignment tool [(4)](https://sciwheel.com/work/citation?ids=48641&pre=&suf=&sa=0) within the in-house DKFZ One Touch Pipeline (OTP) of the ODCF. For the identification of Single-Nucleotide Variants (SNVs), our in-house workflow based on samtools [(5)](https://sciwheel.com/work/citation?ids=48787&pre=&suf=&sa=0) mpileup and bcftools [(6)](https://sciwheel.com/work/citation?ids=396559&pre=&suf=&sa=0) (version 2.2.0) was employed (https://github.com/DKFZ-ODCF/SNVCallingWorkflow). Small insertion/deletions (INDELs) were called using the InDelCallingWorkflow (version 3.1.1), which utilizes Platypus (https://github.com/DKFZ-ODCF/IndelCallingWorkflow). The confidence of INDEL variant calls was evaluated using the Platypus scoring system. To detect Structural Variants (SVs), we utilized the Sophia workflow (version 2.2.3) from the ODCF as the standard pipeline (<https://github.com/DKFZ-ODCF/SophiaWorkflow>).

**Processing Samples without Patient-Matched Germline Data: The "No-control workflow"**

For 158 out of 353 (45%) PDXs in our cohort, a corresponding germline control was not available. To address this, we employed an established in-house pipeline "No-Control workflow". To process the PDX and tumor data samples, various tools for variant calling were incorporated: SNVs were called using mpileup, indels were called using Platypus, and SVs were processed using Sophia.

To facilitate the analysis, we soft-linked the aligned tumor BAM files and generated pseudo-control BAM files. The Roddy framework (version 3.5.8) was employed to perform the variant calling and filtering steps (available at https://github.com/TheRoddyWMS/Roddy). To filter out germline variants, we excluded variants that matched entries in the dbSNP and 1K genomes databases.

For variant interpretation and prioritization in both coding and non-coding regions, we utilized the Ensembl Variant Effect Predictor (VEP) [(7)](https://sciwheel.com/work/citation?ids=2708181&pre=&suf=&sa=0). Additionally, to prioritize and include relevant variants, we incorporated gnomAD (version 2.1) [(8)](https://sciwheel.com/work/citation?ids=15743276&pre=&suf=&sa=0), a widely used database providing minor allele frequency (MAF) information across various populations. Variants with a maximum population allele frequency exceeding 0.001 were excluded from further consideration as they were likely to represent common polymorphisms. This filtering step enabled us to focus on variants that were less prevalent in the general population, allowing us to identify variants with higher rarity or specific enrichment patterns within our cohort. This approach enhanced the identification of potentially pathogenic or functionally significant variants in our analysis.

**DNA Methylation Array Profiling**

Genomic DNA extracted from patient tumor and PDX samples (fresh frozen material) was subjected to methylation profiling at the DKFZ Genomics and Proteomics Core Facility (Heidelberg, Germany) and Institut Curie using the Methylation450K and MethylationEPIC BeadChip arrays (Illumina). DNA methylation data were analyzed using the “RnBeads” Bioconductor package [(9)](https://sciwheel.com/work/citation?ids=6833361&pre=&suf=&sa=0). Briefly, data import was performed using “rnb.run.import” function. Subsequently, QC analysis was performed, and samples with outlier intensities in 450k/EPIC array control probes were removed from the dataset (https://www.bioconductor.org/packages/release/bioc/vignettes/RnBeads/inst/doc/RnBeads.pdf). This was followed by RnBeads pre-processing module, in which background signal subtraction (“enmix.oob” method, [(10)](https://sciwheel.com/work/citation?ids=1489307&pre=&suf=&sa=0)) and beta-mixture quantile normalization [(11)](https://sciwheel.com/work/citation?ids=506635&pre=&suf=&sa=0) were applied. Unreliable probes (Greedycut algorithm with detection P > 0.01), probes with low standard deviation across samples (<0.005), probes with missing values for > 50% samples, SNP-containing probes, cross-reactive probes, and probes mapping to sex chromosomes were removed. Similarly, unreliable samples were filtered out using Greedycut approach. Samples from 450k and EPIC arrays were pre-processed separately, and overlapping probes were subsequently merged to get a final methylation object. Tumor class annotation was performed using the “brain v12.8” [(12,13)](https://sciwheel.com/work/citation?ids=4946086,18302236&pre=&pre=&suf=&suf=&sa=0,0) and “sarcoma v 13” [(14,15)](https://sciwheel.com/work/citation?ids=10360173,18302237&pre=&pre=&suf=&suf=&sa=0,0) DNA methylation-based tumor classifiers, along with unsupervised clustering analyses using the t-distributed stochastic neighbor embedding (t-SNE) method. By leveraging these classifiers and machine learning techniques, we could predict the tumor type and molecular subtype for each analyzed sample. This approach incorporated external tumor reference datasets [(12,14,16–25)](https://sciwheel.com/work/citation?ids=15418036,8685657,5256095,1006277,9351894,4946086,10360173,3874900,7084515,6822914,12969551,1557222&pre=&pre=&pre=&pre=&pre=&pre=&pre=&pre=&pre=&pre=&pre=&pre=&suf=&suf=&suf=&suf=&suf=&suf=&suf=&suf=&suf=&suf=&suf=&suf=&sa=0,0,0,0,0,0,0,0,0,0,0,0&dbf=0&dbf=0&dbf=0&dbf=0&dbf=0&dbf=0&dbf=0&dbf=0&dbf=0&dbf=0&dbf=0&dbf=0) (**Supplementary Table 2**), leading to improved accuracy and classification reliability. Whenever the molecular subtype of PDX models could not be properly annotated by either the classifier or the clustering methods, models were classified using the following nomenclature: “tumor type (e.g HGG, NB)_nd”. To assess molecular similarity in paired tumor-PDX samples, beta values for all SNP probes were extracted, and pairwise Manhattan distances (M) were calculated (scale 0-1). An arbitrary threshold of M> 0.1 was used to identify potentially divergent paired samples. Purity scores based on methylation array data were inferred using the RF_purify method [(26)](https://sciwheel.com/work/citation?ids=12086937&pre=&suf=&sa=0).

**Copy number variant (CNV) workflow**

To ensure precise identification and characterization of copy number variants (CNVs), multiple bioinformatics tools were used, taking into consideration their individual attributes such as specificity, sensitivity and confidence scores.

The ichorCNA tool [(27)](https://sciwheel.com/work/citation?ids=4477601&pre=&suf=&sa=0), uses a probabilistic Hidden Markov Model (HMM) to segment the genome and predict large-scale copy number alterations from ultra-low-pass whole genome and whole exome data also, in samples with low tumor purity. It employs a deconvolution algorithm to estimate tumor purity and ploidy. For samples without controls, we generated a panel of normal controls using existing controls to reduce noise and improve the accuracy of CNV calling. The output ploidy, tumor purity and copy number alterations were interpreted for further analysis.

Sequenza [(28)](https://sciwheel.com/work/citation?ids=3797373&pre=&suf=&sa=0) is a computational tool designed for the analysis of cancer genomic data. It employs a probabilistic model that specifically operates on paired tumor-normal DNA sequencing data to calculate copy-number profiles, estimate tumor cell fraction, and determine tumor ploidy using whole exome and genome data. To run Sequenza, the required input parameters included aligned BAM files, a human-mouse hybrid genome (hs37d5-GRCm38mm10), and the human reference genome (hs37d5). For whole exome sequencing (WES) data, a bin size of 100 bases was utilized. We used the Sequenza output to accurately estimate tumor purity between patient tumors and PDX models and also enable reliable copy number calls.

CNVkit [(29)](https://sciwheel.com/work/citation?ids=2016124&pre=&suf=&sa=0) (version 0.9.3) is a tool specifically designed to detect and quantify genomic amplifications and deletions in tumor samples. It utilizes a target capture-based sequencing approach to analyze read depths and calculate copy number profiles across the genome. Using the "in-house" standard pipeline with default parameter settings on whole exome sequencing data of PDX and tumor samples, only samples with germline controls could be analyzed using this tool. The log_2_ ratio values from the processed CNVkit output were extracted and concatenated for each fragment and all the ITCC-P4 samples. The results of these analyses were used for comparative analysis of tumor cell purity and to annotate the copy number alterations of driver genes. TCF values were obtained from CNVkit results for samples with available CNVkit outputs.

Copy-number variations (CNVs) were also inferred from DNA methylation data with the conumee package (version 1.9.0) and visualized using IGV (version 2.13; Broad Institute).

**RNA Sequencing Workflow and Gene Fusion Identification**

RNAseq was performed for samples using the HiSeq 2000 instrument (Illumina) at the collaborating partner sites. RNA sequencing data underwent processing using the DKFZ OTP in-house pipeline (version 3.0.0) provided at https://github.com/DKFZ-ODCF/RNAseqWorkflow, which is based on the well-established Roddy workflow (https://github.com/TheRoddyWMS/Roddy). The reads obtained from the samples were aligned to the human reference genome (hg37d5) employing the STAR aligner [(30)](https://sciwheel.com/work/citation?ids=49324&pre=&suf=&sa=0). Subsequently, the mapped BAM files were utilized in FeatureCounts [(31)](https://sciwheel.com/work/citation?ids=148598&pre=&suf=&sa=0) to quantify the gene expression using the GENCODE v19 reference annotation. To identify gene fusions, the input RNA-sequencing data were subjected to analysis using Arriba (version 0.8) [(32)](https://sciwheel.com/work/citation?ids=10311000&pre=&suf=&sa=0). Only gene fusions reported by Arriba with high or medium confidence levels were extracted and included in further analysis. Expression data were imported in the R2 portal for downstream (unsupervised clustering, differential gene expression) analyses. Gene ontology analyses were performed using the “ShinyGO 0.77” web tool (http://bioinformatics.sdstate.edu/go/).

**Identification and Annotation of Driver Genes for Mutational Landscape Analysis**

We aimed to identify and annotate important driver genes for the analysis of the mutational landscape. To accomplish this, we relied on two key references: the 2022 World Health Organization (WHO) Classification of Pediatric Tumors [(33)](https://sciwheel.com/work/citation?ids=12184736&pre=&suf=&sa=0) and a published review by Jones et al. focusing on pediatric solid cancers [(34)](https://sciwheel.com/work/citation?ids=7192150&pre=&suf=&sa=0) to curate a comprehensive list of known somatic and germline driver mutations.

**Analysis of Tumor Mutational Burden (TMB)**

Tumor mutational burden (TMB) [(35)](https://sciwheel.com/work/citation?ids=11322282&pre=&suf=&sa=0&dbf=0) was calculated for the ITCC-P4 samples based on single nucleotide variants (SNVs), and small insertions/deletions (indels) determined by whole-exome sequencing (WES) data, focusing on the coding regions targeted by the respective WES library preparation kits. Only high-confidence, somatic and functional exonic SNVs and indels were considered for TMB calculation. The obtained results were compiled and utilized for comparing the mutation load across different cancer subgroups, disease states, and model types.

**Calculation of Variant Allele Frequency**

To gain insights into intra-tumor heterogeneity, clonal selection, the distinction between somatic and germline variants, and the evolution within the PDX samples, we performed calculations to determine the variant allele frequency (VAF) of functional somatic SNVs. The first step involved extracting the DP4 values from the aligned and processed somatic functional SNV files. The DP4 value represents the sequencing depth for each allele (reference and alternate) in each sample and is commonly used to indicate the number of reads supporting each allele. It comprises four subfields that indicate the coverage of the reference allele by forward reads, the coverage of the reference allele by reverse reads, the coverage of the alternate allele by forward reads, and the coverage of the alternate allele by reverse reads. Subsequently, the VAF scores were calculated using the following formula:

$$VAF=(Forward non-ref allele+Reverse non-ref allele)/(Forward ref +Reverse ref+Forward non-ref+Reverse non-ref)$$

**PDX-Tumor correlation analysis using delta-VAF and VAF-ratios**

In order to assess and establish the correlation of clonal discordance between the variant allele frequencies of tumors and PDX models, we introduced a metric called “deltaVAF”. This metric was derived by subtracting the tumor VAF from the PDX VAF. The identification of PDX-specific variants within a sample played a crucial role in the investigation of significant subclones. To analyze the presence of overlapping or exclusive variants between PDX and tumor, we introduced another metric known as "VAFratio." The VAFratio scores were calculated by summing the PDX VAF and dividing it by the sum of the tumor VAF for each patient sample, facilitating a quantitative comparison of variant frequencies.

**Statistical analysis**

All statistical calculations (such as Pearson correlation and log2 fold-change values) and downstream molecular analyses for this study were conducted, and graphical visualizations were generated within the R environment.
